# Pdk1-dependent metabolic heterogeneity promotes aggressive colorectal cancer through cell-intrinsic interferon signaling

**DOI:** 10.64898/2026.10.07.757309

**Authors:** Jessica Calviello, Lisa Pöltl, Alessia Mira, Filippo Spini, Ainhoa Larrañaga, Lídia Fortuny, Federica Vianello, Andrea Bonilla-Díaz, Albert Álvarez-Muñoz, Giulia Pazzagli, Maria Serra, Nadia Ducano, Marta Gut, Eloi Casals, Anna Esteve-Codina, Marc Liesa, Livio Trusolino, Anna Sapino, David Sebastián, Carlos Sebastián

**Author notes:** These authors equally contributed to this work (Jessica Calviello, Lisa Pöltl). Department of Molecular Biotechnology and Health Sciences, Molecular Biotechnology Center, University of Torino; Torino, Italy (Alessia Mira). Department of Biology, University of Padova; 35131 Padova, Italy (Federica Vianello).

## Abstract

Tumor heterogeneity is a major determinant of tumor progression and therapeutic resistance, yet the contribution of metabolism to intratumoral heterogeneity still remains poorly explored. Here, we find that pyruvate dehydrogenase kinase-1 (Pdk1) expression defines metabolically distinct cancer cell populations in colorectal tumors. Image-based spatial single-cell phenotyping shows that Pdk1^high^ cells are associated with tumor progression in orthotopic models of colorectal cancer (CRC). Genetic ablation experiments demonstrate the role of Pdk1^high^ cells in tumor growth, 5-fluorouracil chemoresistance and tumor relapse. Mechanistically, we identify a link between Pdk1-driven metabolic states and specific cell phenotypes, where Pdk1^high^ cancer cells exhibit an EMT, stem cell-like phenotype characterized by increased interferon (IFN) signaling. Mitochondrial remodeling in Pdk1^high^ cells leads to the release of mitochondrial DNA to the cytosol and activation of IFN pathways, promoting Pdk1-driven phenotypes. Inhibition of IFN signaling impairs the tumorigenic and aggressive features of Pdk1^high^ cancer cells, thus establishing a functional connection between metabolic heterogeneity, cell-intrinsic IFN signaling and aggressive CRC.

## Introduction

Despite recent advances in the diagnosis and treatment of colorectal cancer (CRC), this cancer type is the third most common malignancy worldwide and the second leading cause of cancer-related deaths (*1*). The disease becomes challenging to treat at advanced stages and eventually develops resistance to most forms of combination therapy. Intratumoral heterogeneity (ITH), defined as the presence of diverse subpopulations of cancer cells within a tumor, is a major contributor to tumor progression and relapse in a variety of tumor types, including CRC (*2, 3*). Both genetic and non-genetic mechanisms have been described to promote ITH. Genetic heterogeneity refers to the existence of subclonal populations of cancer cells with different genotypes within a tumor, mainly caused by somatic mutations conferring a selective growth advantage. Non-genetic ITH, on the other hand, involves epigenetic, transcriptional and translational heterogeneity leading to the acquisition of different cell states and phenotypes. These phenotypic changes are plastic and evolve during tumor progression, mainly due to microenvironmental cues, making cancer cell plasticity a major contributor of ITH, tumor progression and resistance to therapies (*4*).

Metabolic reprogramming is a key feature of cancer cells. Rewiring of core metabolic pathways allows cancer cells to adapt to the microenvironmental conditions of a growing tumor, thus promoting tumor progression (*5*). However, the contribution of cancer cell metabolism to ITH has been less explored. Few studies have reported the presence of intratumoral metabolic heterogeneity in several tumor types (*6–11*), which has been mainly associated to microenvironmental factors, such as proximity to blood vessels, hypoxia, nutrient availability and the interaction of cancer cells with stromal and immune cells present in the tumor microenvironment (TME)(*12*). However, our understanding of the metabolic complexity of tumors, its dynamic regulation during tumor progression and its biological consequences on cancer evolution and response to therapies is still very limited. Furthermore, whether cell-intrinsic metabolic features could determine specific cell phenotypes contributing to ITH remains poorly understood. Our recent work identified a rare population of highly glycolytic cells within the intestinal epithelium and *APC^nin^* adenomas characterized by high levels of pyruvate dehydrogenase kinase (Pdk)-1 (*13*). However, whether this cell population drives a similar metabolic compartmentalization in CRC and its role on tumor progression remained to be elucidated. Here, we found that Pdk1 expression establishes an intratumor metabolic heterogeneity in CRC, which functionally contributes to tumor progression, therapy resistance and tumor relapse. Our results show that Pdk1^high^ cancer cells exhibit an epithelial-to-mesenchymal transition (EMT) and a stem cell-like phenotype, characterized by cell-intrinsic activation of interferon (IFN) pathways. Importantly, we found that the metabolic state of Pdk1^high^ cancer cells links mitochondrial remodeling to the release of mitochondrial nucleic acids to the cytosol and activation of IFN signaling in these cells, contributing to tumor growth. Our results provide insights into the metabolic regulation of tumor heterogeneity and its contribution to CRC aggressiveness, identifying the Pdk1-IFN axis as a potential therapeutic target.

## Results

### Visualizing Pdk1-dependent metabolic heterogeneity in CRC

Our previous work identified Pdk1 activity as a marker of a subpopulation of highly glycolytic cells within the intestinal epithelium and adenomas (*13*). To further characterize this metabolic heterogeneity and the presence of Pdk1^high^ cells in CRC, we took advantage of our recently described *Pdk1*-mCherry reporter, which allows the direct visualization of single cells with different levels of *Pdk1* expression (*13*). We used MC38 cells, a CRC cell line derived from C57BL6/J mice, to generate MC38-*Pdk1*-mCherry cells. Analysis of subcutaneous tumors derived from these cells revealed a heterogenous pattern of Pdk1 expression (fig.S1A), in agreement with our previous work (*13*). Next, we analyzed the expression of the *Pdk1*-mCherry reporter in the context of colorectal tumors growing in their native microenvironment by transplanting MC38-*Pdk1*-mCherry cells in the caecum of C57BL6/J mice. Remarkably, analysis of histological sections from these orthotopic tumors showed the presence of Pdk1^high^ and Pdk1^low^ cancer cells throughout the tumor (Fig.1A). Similar results were obtained in subcutaneous and orthotopic tumors derived from the human CRC cell line HCT116 carrying a *PDK1*-mCherry reporter (fig. S1B-C). To obtain a 3D picture of Pdk1 expression in colorectal tumors, we performed large-scale single-cell resolution 3D (LSR-3D) imaging, a recently described method to visualize normal tissue architecture and entire tumors at cellular resolution (*14*). Our results confirmed Pdk1-dependent spatial metabolic heterogeneity at 3D level in both subcutaneous and orthotopic tumors derived from MC38-*Pdk1*-mCherry cells as depicted by the heterogeneous expression of the reporter (Fig.1B and fig.S1D), demonstrating that colorectal tumors exhibit Pdk1-dependent glucose metabolism heterogeneity.

**Figure 1.**
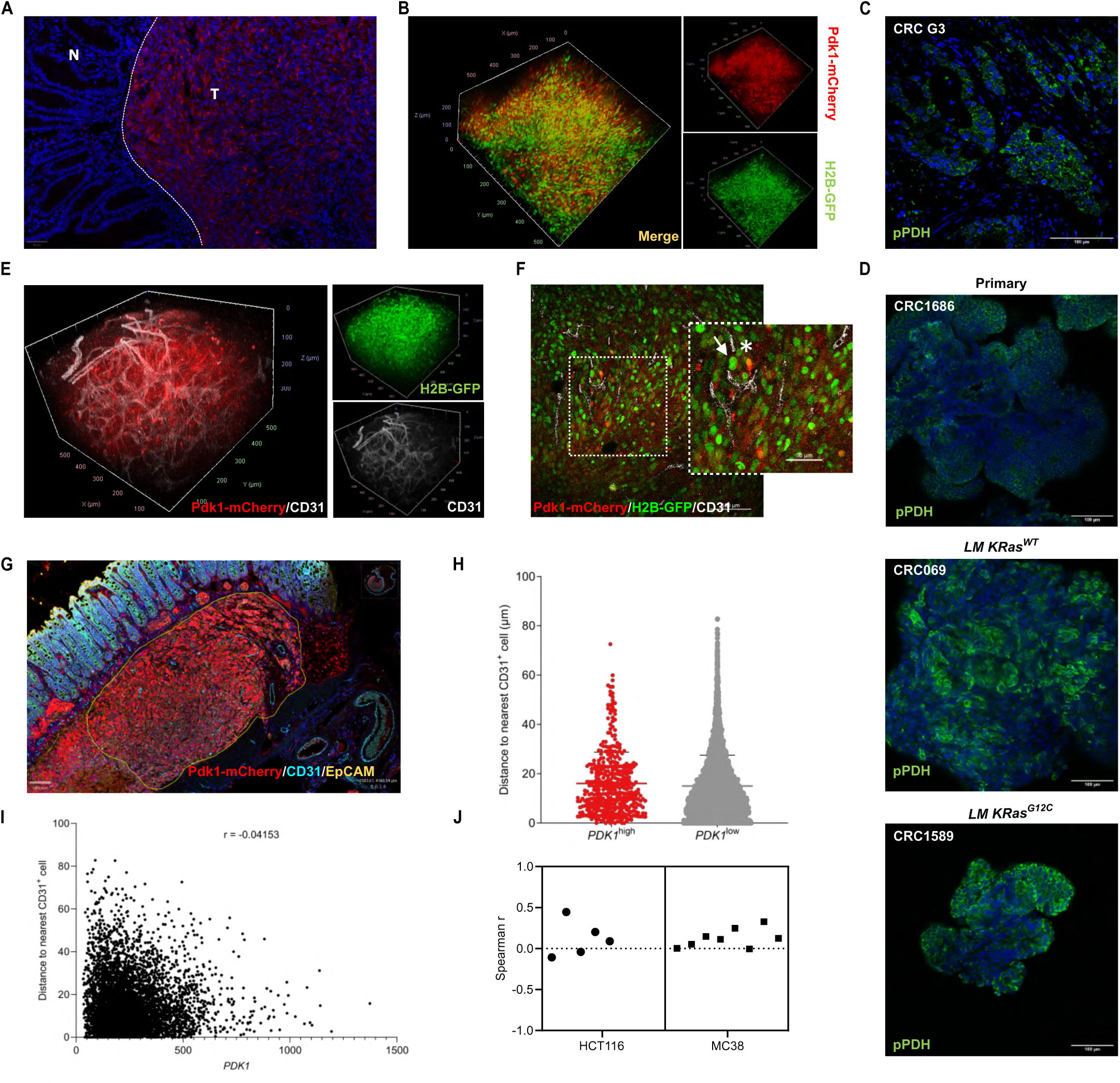
Pdk1-dependent metabolic heterogeneity in CRC. **(A)** Representative fluorescence microscopy image of an orthotopic colorectal tumor section derived from MC38-Pdk1-mCherry cells. Scale bar, 50μm. **(B)** Representative LSR-3D image of a subcutaneous tumor derived from MC38-Pdk1-mCherry cells stably expressing H2B-GFP. **(C)** pPDH staining of a section from a G3 human colorectal tumor. Scale bar, 100μm. **(D)** pPDH staining of PDOs derived from primary CRC (upper panel) and liver metastatic lesions *KRas^WT^*(middle panel) and *KRas^G12C^* (lower panel). **(E)** LSR-3D imaging of a subcutaneous tumor derived from the cells in B and stained with anti-CD31 to visualize blood vessels. **(F)** Representative fluorescence microscopy image of a section from tumor in E. A magnified inset shows Pdk1-mCherry^high^ cells located in close proximity to CD31^+^ blood vessels (white asterisk) and Pdk1-mCherry^low^ cells in similar perivascular regions (white arrow). Scale bar, 100μm; inset scale bar, 50μm. **(G)** Representative multiplex immunofluorescence image of an orthotopic tumor section derived from HCT116-PDK1-mCherry cells, stained for mCherry (red), CD31 (cyan, blood vessels), EpCAM (yellow, epithelial cells) and DAPI (nuclei). The yellow outline delimits the tumor mass considered for analysis. Cyan staining of normal intestinal epithelium is considered to be non-specific. Scale bar, 100 μm. **(H)** Violin plot showing the distribution of distances (μm) to the nearest CD31^+^ endothelial cell for PDK1^high^ and PDK1^low^ cancer cells, quantified by spatial single-cell phenotyping using QuPath software. **(I)** Scatter plot showing the correlation between PDK1 fluorescence intensity (x-axis) and distance to the nearest CD31^+^ cell (y-axis) for individual cancer cells within the tumor. The Pearson correlation coefficient (r = -0.04) confirms the absence of any meaningful correlation between PDK1 expression level and vascular proximity. **(J)** Summary scatter plot showing the correlation between PDK1 fluorescence intensity and distance to the nearest CD31^+^ endothelial cell across all tumor regions analyzed in both HCT116 (n=5) and MC38 (n=8) orthotopic tumors.

To extend this observation to human colorectal tumors and validate PDK1 expression heterogeneity in human CRC, we stained tumor sections form CRC patients with anti-phospho-PDH (Ser293) as a readout of PDK1 activity, which we have previously shown to mark Pdk1^high^ cells (*13*). Consistent with our results above, we found pPDH^+^ and pPDH^-^ cells in high grade (G3) colorectal carcinomas (Fig. 1C), as well as in adenomas and low/moderate grade (G1/G2) CRC (fig. S1E), suggesting that metabolic heterogeneity is established early during tumorigenesis. We also performed pPDH staining in the XENTURION collection of patient-derived organoids (PDOs)(*15*), including 5 PDO lines derived from primary colorectal tumors, 5 PDO lines derived from *KRas^WT^* liver metastases and 5 PDO lines derived from liver metastases harboring mutations in *KRas*. Again, our results showed a heterogeneous pattern of pPDH staining in all the PDO lines analyzed (Fig. 1D and fig. S1F). Altogether, these results indicate that colorectal tumors are formed by cancer cells with different metabolic activities, revealing the presence of intratumoral glucose metabolism heterogeneity in CRC.

### Spatial analysis of Pdk1^high^ cells during tumor progression

Oxygen and nutrient gradients, which are influenced by distance to blood vessels, have been characterized as main determinants of metabolic heterogeneity in tumors (*12*). Moreover, stabilization of Hif1α by hypoxia has been shown to induce the expression of *Pdk1* (*16*). To analyze the spatial distribution of Pdk1^high^ colorectal cancer cells and whether their metabolic phenotype depends on their distance to blood vessels, we performed LSR-3D imaging of both subcutaneous and orthotopic MC38-*Pdk1*-mCherry tumors in combination with CD31 staining to mark endothelial cells. We found Pdk1^high^ cells to be distributed throughout the tumor, even next to blood vessels (Fig. 1E, 1F and fig. S2A, S2B), suggesting that Pdk1 expression and the observed metabolic heterogeneity is not linked to differential nutrient availability or hypoxia. To quantitatively analyze the relative distances between Pdk1^high^/Pdk1^low^ cells and blood vessels and to shed light on the contribution of Pdk1^high^ cells to tumor progression, we performed multiplex immunofluorescence experiments coupled with spatial single-cell phenotyping. We stained orthotopic tumors derived from HCT116-*PDK1*-mCherry cells with antibodies detecting mCherry, EpCam (epithelial cell marker) and CD31 (Fig. 1G). Distance measurements between PDK1^high^/PDK1^low^ cancer cells and CD31^+^ endothelial cells showed no differences in the spatial distribution of these two cell subpopulations of cancer cells relative to blood vessels (Fig. 1H, 1I). We also confirmed these results in MC38-*Pdk1*-mCherry orthotopic tumors. Interestingly, we found some tumor areas where the average distance of Pdk1^high^ cells to CD31^+^ cells was slightly higher compared to Pdk1^low^ cells, while other tumor areas showed the opposite phenotype (fig. S2C-H). However, there was no correlation between mCherry fluorescence and the distance to blood vessels in any of the regions analyzed in both models (Fig. 1J), indicating that, overall, the metabolic phenotype of Pdk1^high^ cells is not imposed by oxygen/nutrient gradients.

The use of orthotopic tumors allowed us to identify histopathological areas within these tumors with different aggressive features, including the tumor core (TC), invasive fronts (IF) and tumor buds/poorly-differentiated clusters (PDCs) (Fig. 2A). Our single-cell phenotyping analysis showed that the tumor core contained around 5% of PDK1^high^ cells, and most of these cells were slow cycling/quiescent (Ki67^-^) (Fig. 2B and 2C), in agreement with our previous findings (*13*). Remarkably, the percentage of PDK1^high^ cells increased in IFs and PDCs, reaching around 40% in the latter (Fig. 2B). These morphological features have been associated with infiltrative and aggressive growth and are strong negative prognostic factors in CRC (*17*), suggesting that highly glycolytic PDK1^high^ cancer cells could be involved in tumor progression. Interestingly, this expansion in the number of PDK1^high^ cells in IFs and PDCs was associated with an increase in PDK1^high^Ki67^+^ double positive cells (Fig. 2C), indicating a phenotypic switch of PDK1^high^ cells from quiescence to proliferation.

**Figure 2.**
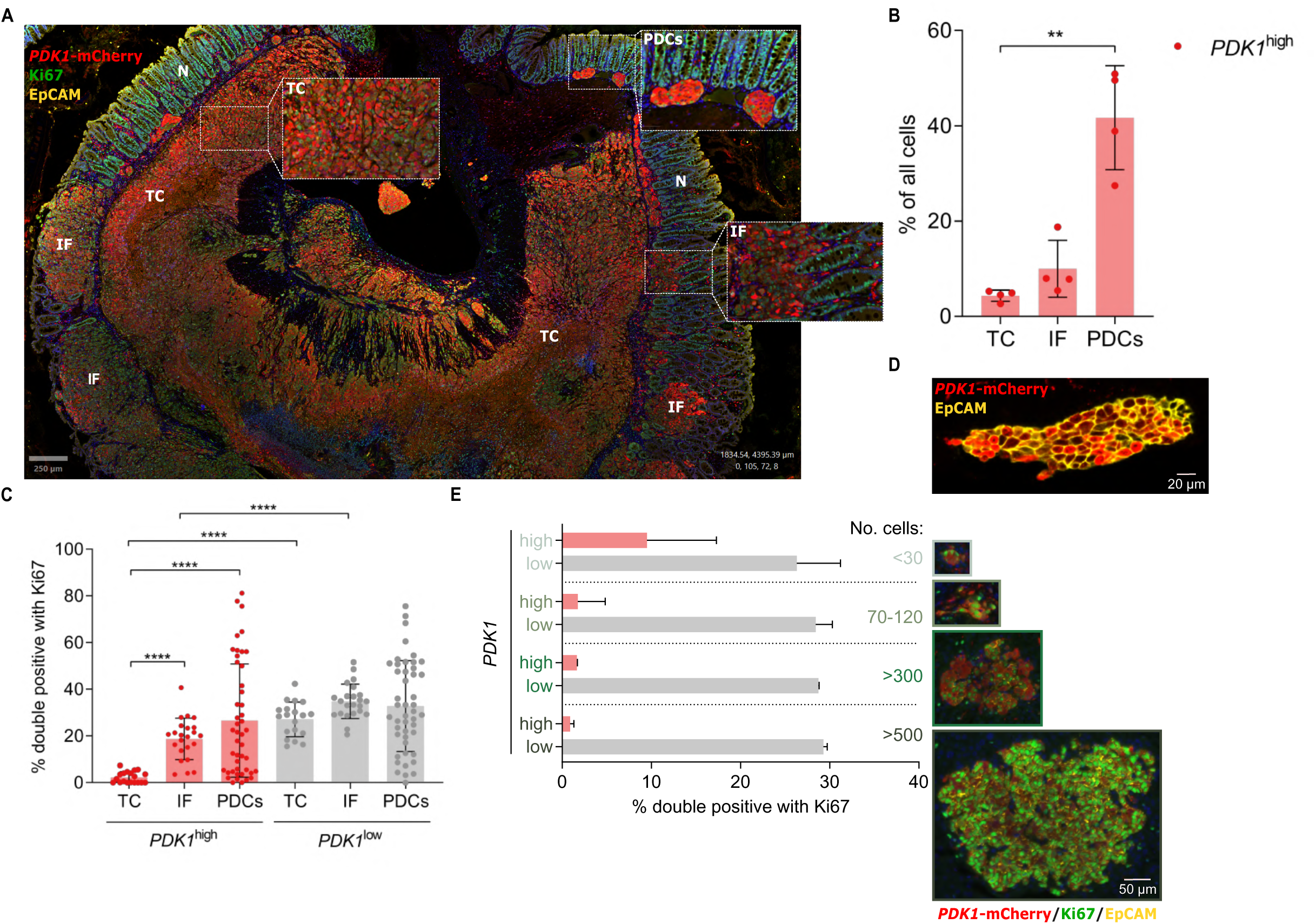
Spatial analysis of Pdk1^high^ cells during tumor progression. **(A)** Immunofluorescent image of an orthotopic cecal tumor marking PDK1-mCherry (red), Ki67 (green) and EpCAM (yellow) reveals metabolically heterogeneous PDK1-expression in different histopathological areas. N = normal; TC = tumor core; IF = invasive front; PDCs = poorly differentiated clusters. Scale bar, 250 μm **(B)** Percentage of PDK1^high^ cells in TC, IF and PDCs. Each data point represents the sum of cells analyzed from different ROIs of n=4 mice. Total cells analyzed: TC: 92745, IF: 20043, PDCs: 5913. Statistical significance was assessed using Kruskal-Wallis test (p**<0.01). **(C)** Percentage of Ki67^+^ PDK1^high^ and PDK1^low^ cells in TC, IF and PDCs. Each data point represents an ROI from n=5 mice (total 110539 cells) for TC, n=6 mice (total of 31837 cells) for IF and n=5 mice (total of 8099 cells) for PDCs. Statistical significance was assessed using Kruskal-Wallis test (between areas) and Mann-Whitney test (within areas) (p****<0.0001). **(D)** Representative multiplex immunofluorescence image of a liver metastatic lesion derived from HCT116-PDK1-mCherry cells, stained for mCherry and EpCAM. Scale bar, 20 μm. **(E)** Quantification of the percentage of PDK1^high^ Ki67^+^ and PDK1^low^ Ki67^+^ cells stratified by metastasis size, defined by the number of cells per metastatic lesion: micrometastases (<30 cells, n=5), small metastases (70–120 cells, n=3), medium metastases (>300 cells, n=2), and large metastases (>500 cells, n=2). Representative fluorescence images of a metastatic lesions from each size category are shown alongside the quantification. Data are expressed as mean ± SD. Scale bar, 50 μm.

Next, we sought to investigate whether PDK1^high^ cells are present in CRC metastases. Thus, we expanded our analysis to metastatic lesions in the liver and observed that they contained both PDK1^high^ and PDK1^low^ cells (Fig. 2D), thus recapitulating the metabolic heterogeneity observed in primary tumors. Of note, 10% of PDK1^high^ cells were Ki67^+^ in micrometastases, while the number of proliferating PDK1^high^ cells tended to decrease as the metastases grew bigger (Fig. 2E). These results suggest that PDK1^high^ cells could be important for the establishment of the metastatic lesion, whereas proliferative PDK1^low^ cells drive metastatic growth. Taken together, our results demonstrate the presence of PDK1-driven spatiotemporal metabolic heterogeneity in CRC and suggest a key role for PDK1^high^ cells in tumor progression.

### Functional analysis of Pdk1^high^ colorectal cancer cells

Based on our results above, we reasoned that Pdk1-dependent metabolic heterogeneity could be a key contributor to tumor progression. To test this, we developed a genetic ablation system to eliminate Pdk1^high^ cells. We generated MC38 cells carrying *Pdk1*-CreERT2 and flexDTA reporters, which allow the selective ablation of Pdk1-expressing cells by 4-hydroxytamoxifen (4-OTH)-induced and Cre-mediated expression of Diphtheria toxin A (*18*). To visualize Pdk1^high^ and Pdk1^low^ cells, we also transduced these cells with the *Pdk1*-mCherry reporter, generating MC38-*Pdk1*-mCherry/CreERT2-flexDTA cells (MC38-*Pdk1*-mCh/Cre-DTA, Fig.3A). These cells were first used to generate 3D-spheroids, an in vitro system that better mimics the complexity and heterogeneity of tumors, as they allow cell-to-cell and cell-to-ECM interactions as well as the establishment of metabolic gradients (*19*). In line with this, we observed a heterogeneous pattern of quiescent Pdk1^high^ cells in MC38-*Pdk1*-mCh/Cre-DTA spheroids (Fig.3B), consistent with our results in tumors. Addition of 1μM of 4-OHT reduced the number of mCherry^high^ cells by 50% without barely affecting mCherry^low^ or mCherry^med^ cells (fig. S3A). In these conditions, ablation of Pdk1^high^ cells impaired spheroid growth (Fig. 3C and 3D). Similar results were obtained using HCT116-*PDK1*-mCh/Cre-DTA cells (fig. S3B). To rule out a non-specific toxic effect of 4-OHT in Cre-expressing cells, we performed the same experiment in spheroids formed by MC38-*Pdk1*-mCh/Cre cells (not expressing the DTA cassette) and found no differences in spheroid size (fig. S3C). Furthermore, we confirmed that this phenotype was dependent on the selective elimination of Pdk1^high^ cells, as 4-OHT treatment did not reduce the growth of Pdk1-mCherry^low^ cells-derived spheroids while completely impaired spheroid formation from Pdk1-mCherry^high^ sorted cells (fig.S3D).

**Figure 3.**
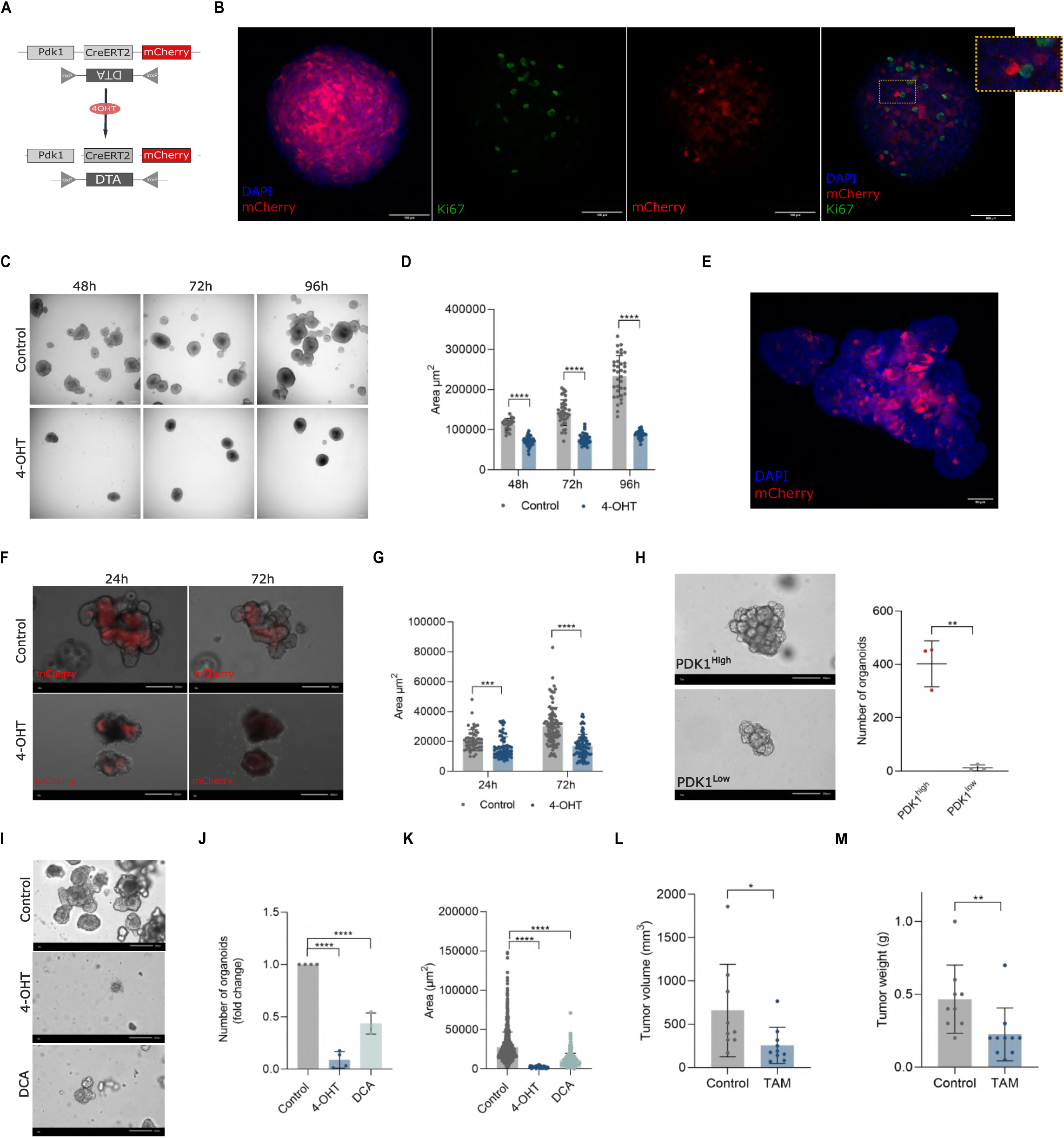
Functional analysis of Pdk1^high^ colorectal cancer cells. **(A)** Schematic representation of the Pdk1-CreERT2/FlexDTA genetic ablation system. **(B)** Representative confocal Z-stack image of an MC38-Pdk1-mCherry spheroid showing the heterogeneous distribution of Pdk1-mCherry signal Ki67 expression. Magnified inset highlights the absence of overlap between individual Pdk1-mCherry^high^ cells and Ki67^+^ nuclei. **(C)** Representative brightfield images of MC38-Pdk1-mCherry/CreERT2-FlexDTA spheroids treated with vehicle (control) or 4-OHT at 48-, 72-, and 96-hours post-treatment. **(D)** Quantification of spheroid area (µm²) at 48, 72, and 96 hours in vehicle-treated (control) and 4-OHT-treated spheroids. Data shown are from a representative experiment (n = 10). Data are expressed as mean ± SD. Statistical significance was assessed using multiple t-test, ****p < 0.0001. **(E)** Confocal microscopy image of CRC1676 PDOs expressing the PDK1-mCherry reporter stained with DAPI (nuclei), acquired at 20× magnification. Scale bar, 50μm. **(F)** Representative brightfield and fluorescence images of the same CRC1676 organoids tracked over time, showing vehicle (control) or 4-OHT treated organoids at 24- and 72-hours post-treatment. Scale bar, 200μm. **(G)** Quantification of organoid area (µm²) in CRC1676 PDOs treated with vehicle (control) or 4-OHT at 24- and 72-hours post-treatment. A representative experiment is shown (n=4). Statistical significance was assessed using multiple t-test, ***p < 0.001, ****p < 0.0001. **(H)** Representative brightfield images of PDOs grown from single-cell suspensions of FACS-sorted PDK1^high^ and PDK1^low^ cells 15 days after being plated. The right panel shows the quantification of organoid number at day 15 (n = 3). Data are expressed as mean ± SEM. Statistical significance was assessed using multiple t-test, **p < 0.01. Scale bar, 200μm. **(I)** Representative brightfield images at day 15 of organoid formation assay from single-cell suspensions of PDOs treated with vehicle (control), 4-OHT (1 μM), or DCA (10 mM). Scale bar, 200μm. **(J)** Quantification of organoid number at day 15 in organoid formation assays from single-cell suspensions of PDOs treated with vehicle (control), 4-OHT, or DCA (n = 3). Data are expressed as mean ± SEM. Statistical significance was assessed using multiple t-test, ****p < 0.0001 **(K)** Quantification of organoid area (µm²) at day 15 in organoid formation assays from single-cell suspensions of PDOs treated with vehicle (control), 4-OHT, or DCA (n = 3). Data are expressed as mean ± SD. Statistical significance was assessed using multiple t-test. ****p < 0.0001. **(L)** Quantification of tumor volume (mm³) at experimental endpoint (day 20) of subcutaneous MC38-Pdk1-mCherry/CreERT2-FlexDTA tumors treated with vehicle (control) or tamoxifen (TAM; 2 mg per injection, every two days) (n = 9). **(M)** Quantification of tumor weight (g) at experimental endpoint (day 20) (n = 9). Data are expressed as mean ± SD. Statistical significance was assessed using Mann-Whitney test. *p < 0.05, **p < 0.01.

To extend these experiments to a more relevant model of human CRC, we generated three independent CRC PDO lines (CRC1676 and CRC1898, derived from primary tumors, and CRC1589, derived from a liver metastasis) carrying the *PDK1*-mCh/Cre/DTA system. Similar to MC38 and HCT116 cells, quiescent PDK1^high^ cells were heterogeneously distributed in these PDOs (Fig.3E and fig.S3E). Importantly, ablation of PDK1^high^ cells impaired the growth of PDOs (Fig.3F and 3G), thus confirming the functional relevance of PDK1^high^ cells in CRC.

Next, we investigated whether, as observed in normal small intestinal organoids (*13*), PDK1^high^ cells within PDOs exhibited stem cell potential. We sorted PDK1-mCherry^high^ and PDK1-mCherry^low^ cells from PDOs and performed organoid formation experiments from sorted single cells. Our results showed increased organoid formation efficiency of PDK1-mCherry^high^ cells (Fig.3H), a readout of stem cell/tumorigenic potential. Moreover, addition of 4-OHT during organoid formation from unsorted single cells almost completely impaired organoid growth compared to control conditions (Fig. 3I-K), suggesting that most newly-formed organoids arise from PDK1^high^ cells. Remarkably, similar results were obtained with the PDK1 inhibitor dichloroacetate (DCA) (Fig. 3I-K), indicating that PDK activity is required for the stem cell/tumorigenic potential of PDK1^high^ cells.

Then, we assessed whether ablation of Pdk1^high^ cancer cells could have an effect on tumor growth *in vivo*. To do so, we injected MC38-*Pdk1*-mCh/Cre-DTA cells into the flanks of C57BL6/J mice and analyzed tumor size upon tamoxifen administration. In agreement with our *in vitro* experiments, tamoxifen treatment resulted in smaller tumors (Fig.3L and 3M), indicating that Pdk1^high^ cells contribute to tumor growth *in vivo*. Altogether, these results demonstrate that Pdk1^high^ colorectal cancer cells are key regulators of tumor formation and growth, likely due to their stem cell/tumorigenic potential.

### Molecular characterization of the Pdk1^high^ state

Our results indicate that Pdk1 expression defines two functionally distinct cancer cell populations, implying that the cell metabolic status could be associated with particular cell phenotypes. To explore this possibility and to provide insights into the molecular determinants involved in the regulation of tumor progression by Pdk1^high^ cells, we performed comparative transcriptomics by RNA-sequencing using mCherry-sorted cells from MC38-*Pdk1*-mCherry subcutaneous tumors. Our results identified 440 differentially expressed genes (261 genes upregulated, and 179 genes downregulated) in Pdk1^high^ cells compared to Pdk1^low^ cells (fig. S4A and table S1). Gene Set Enrichment Analysis (GSEA) showed downregulation of gene signatures related to cell proliferation, including E2F2-targets, MYC-targets, and DNA replication genes (*Cdt1, Mcm genes, Pcna, Ung, Pola2*) in Pdk1^high^ cells (Fig. 4A-B and fig. S4B-C), in line with their quiescence nature. Indeed, we also found downregulation of genes of metabolic pathways normally associated with cell proliferation, such as nucleoside/nucleotide metabolism (*Cad, Nme1, Shmt, Dhodh, Mthfd1*) and ribosome biosynthesis (*Rrs1, Imp4, Bop1, Rrp9*) (Fig. 4B and fig. S4C). Remarkably, expression of genes involved in mitochondrial structure and organization (*Tomm5, Tomm20, Mrpl3, Timm50*), OxPhos (*Slc25a5, Ndufa4, Atp5b, Ndufa9, Cox4i1*) and TCA cycle (*Mdh2, Fh1, Idh3a*) were also downregulated in Pdk1^high^ cells (Fig. 4B and fig. S4C), indicating a low mitochondrial metabolism state and in agreement with their glycolytic phenotype. Consistent with their metabolic features, we found higher expression of the glycolytic genes *Pdk1*, *Pdk2* and *Hk1* in Pdk1^high^ cells (Fig. S4D).

**Figure 4.**
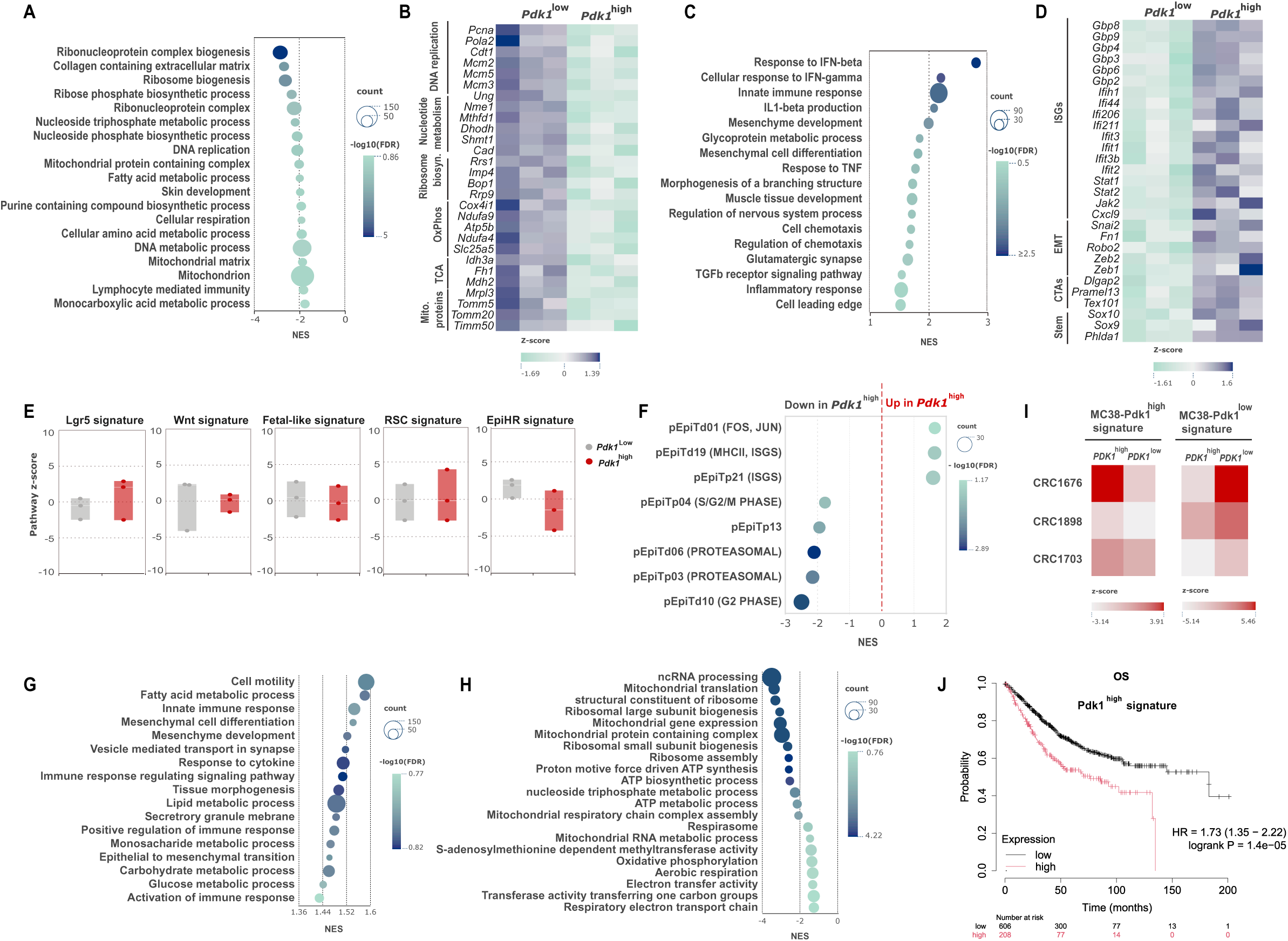
Molecular characterization of the Pdk1^high^ state. **(A)** Bubble plot showing the normalized enrichment scores (NES) of gene ontology (GO) biological process terms significantly downregulated in Pdk1^high^ cells compared to Pdk1^low^ cells sorted form MC38 tumors. Bubble size reflects the number of genes in each gene set and color intensity reflects the -log10(FDR). **(B)** Heatmap showing the row Z-score of selected downregulated genes in Pdk1^high^ cells grouped by biological category. **(C)** Bubble plot showing the normalized enrichment scores (NES) of gene ontology (GO) biological process terms significantly upregulated in Pdk1^high^ cells compared to Pdk1^low^. Bubble size reflects the number of genes in each gene set and color intensity reflects the -log10(FDR). **(D)** Heatmap showing the row Z-score of selected upregulated genes in Pdk1^high^ cells grouped by biological category. **(E)** Box plots showing the pathway enrichment scores of indicated gene signatures in Pdk1^high^ (red) and Pdk1^low^ (grey) cells sorted from MC38-Pdk1-mCherry subcutaneous tumors. **(F)** Bubble plot showing the normalized enrichment scores (NES) of the indicated tumoral epithelial gene signatures from Pelka et al. (*36*) significantly upregulated and downregulated in Pdk1^high^ cells compared to Pdk1^low^ cells. Bubble size reflects the number of genes in each gene set and color intensity reflects the -log10(FDR). Bubble plot showing the NES of gene ontology (GO) biological process terms significantly upregulated **(G)** and downregulated **(H)** in PDK1^high^ cells compared to PDK1^low^ cells sorted from three PDO lines (CRC1676, CRC1898, and CRC1703). Bubble size reflects the number of genes in each gene set and color intensity reflects the - log10(FDR). **(I)** Heatmap showing the enrichment of the MC38-Pdk1^high^ and MC38-Pdk1^low^ gene signatures in PDK1^high^ and PDK1^low^ cells from the three PDO lines (CRC1676, CRC1898, and CRC1703). **(J)** Kaplan-Meier survival analysis of overall survival (OS) in CRC patients stratified by high or low expression of the MC38-Pdk1^high^ gene signature (HR = 1.73, 95% CI 1.35–2.22, log-rank p = 1.4×10⁻⁵). Numbers at risk are shown below the graph.

Among the most enriched gene programs, we found signatures associated with inflammatory responses, type I and type II interferons (IFN) and innate immunity (Fig. 4C and fig. S4B). Many key interferon-stimulated genes (ISGs) were upregulated in Pdk1^high^ cells, including guanylate-binding proteins (*Gbp2, Gbp3, Gbp4, Gbp6, Gbp8, Gbp9),* IFN-induced proteins (Ifih1, Ifi44, Ifi206, Ifi211), IFN-induced protein with tetratricopeptide repeats (Ifit) proteins (*Ifit1, Ifit3, Ifit3b, Ifit2*), signal transducers and transcription factors (*Jak2, Stat1, Stat2*) and chemokines (*Cxcl9*) (Fig. 4D and fig. S4E). Although activation of IFN responses has been usually described as a tumor suppressor mechanism, sustained IFN signaling has been shown to promote tumor progression by inducing stemness and EMT properties in cancer cells (*20–22*). In line with this, we found enrichment of gene signatures related to EMT in Pdk1^high^ cells (mesenchyme development and mesenchymal cell differentiation) (Fig. 4C), which included increased expression of the main EMT genes *Snai2, Zeb2, Zeb1, Fn1* and *Robo2* (Fig. 4D and fig. S4E). Furthermore, among the top upregulated genes in Pdk1^high^ cells, we found tissue-restricted antigens/cancer-testis antigens (*Dlgap2, Pramel13, Tex101*) (Fig. 4D and table S1), suggesting that these cells could be in an undifferentiated state. Consistent with this hypothesis, Pdk1^high^ cells showed upregulation of gene signatures related to differentiation to several tissues, including morphogenesis of a branching structure, muscle tissue development and nervous system development (Fig. 4C), and increased expression of the stem cell/progenitor markers *Sox9*, *Sox10* and *Phlda1* (*23–25*) (Fig. 4D). These results indicate that the Pdk1^high^ state might be linked to stem cell-like and EMT phenotypes, which are compatible with our results showing increased Pdk1 expression in invasive fronts and PDCs (Fig. 2B), enhanced organoid formation efficiency (Fig. 3H) and tumor initiating potential (*13*).

Previous reports have demonstrated that Lgr5^+^ stem-like tumor cells are responsible for CRC growth (*26, 27*). However, pathway enrichment analysis indicated that Pdk1^high^ cells might represent a distinct population with stem cell/tumor initiating potential within colorectal tumors, as no enrichment in Lgr5 and Wnt signatures (*28, 29*) was found in these cells compared to Pdk1^low^ cells (Fig. 4E). Recent studies have identified Lgr5^neg^ cell populations with stem cell-like properties involved in therapy resistance and metastasis formation (*30–34*). These cells are characterized by the activation of regenerative and fetal-like transcriptional programs (*32, 33, 35*), as well as by increased expression of a gene signature associated to high risk of metastatic recurrence (*31*). However, the expression of regenerative stem cells (RSC, fetal-like) and high-risk of relapse (EpiHR) gene signatures were not enriched in Pdk1^high^ cells compared to Pdk1^low^ cells (Fig.4E), indicating that high expression of Pdk1 identifies a distinct cell population or cell state with stem cell-like potential within colorectal tumors.

Next, we sought to analyze whether the Pdk1^high^ signature might be associated to specific cell phenotypes in human CRC. To do so, we first interrogated a recent single-cell RNA-seq dataset comprising 28 mismatch repair proficient (MMRp) and 34 mismatch repair deficient (MMRd) CRC patients, which identified 43 transcriptional programs in malignant cells (*36*). GSEA analysis showed enrichment of immune-related programs from both MMRp and MMRd tumors (ISG, MHC class II and FOS/JUN) in Pdk1^high^ cells, whereas gene programs associated to cell cycle progression (S/G2/M phase) were downregulated (Fig. 4F). These results are in agreement with our observations above and indicate that PDK1^high^ cells might represent a slow cycling/quiescent cell subpopulation with a pro-inflammatory phenotype within human colorectal tumors.

To further characterize the Pdk1^high^ state in human CRC, we took advantage of our collection of PDOs-PDK1-mCherry to perform RNA-seq from mCherry-sorted cells from three independent PDO lines derived from primary tumors (CRC1676, CRC1898, CRC1703). Hierarchical clustering and principal component analysis (PCA) showed that patient of origin is the major grouping factor (fig. S4F), in line with the interpatient heterogeneity observed in CRC. However, analysis of differentially expressed genes identified 297 genes upregulated and 214 genes downregulated in PDK1^high^ cells compared to PDK1^low^ cells (fig. S4G and table S2), indicating that PDK1 expression defines two different cell phenotypes across CRC patients. Indeed, and similar to MC38 tumors, we found enrichment of gene programs related to glycolysis, immune responses (innate immune response, TNFA-signaling, response to cytokine) and mesenchymal cell differentiation in PDK1^high^ cells (Fig. 4G and fig.S4H). Downregulated gene pathways included those associated with cell proliferation (MYC targets, ribosome biosynthesis, nucleotide metabolism) and mitochondrial OxPhos (Fig. 4H and fig. S4H). These results indicate that PDK1^high^ cells within PDOs and MC38 tumors exhibit very similar phenotypic features. Supporting this idea, we found enrichment of the MC38-Pdk1^high^ gene signature and downregulation of the MC38-Pdk1^low^ gene signature in PDK1^high^ cells from PDOs (Fig. 4I). Importantly, the gene signatures defining both MC38-Pdk1^high^ cells and PDOs-PDK1^high^ cells were associated to overall survival of CRC patients, with patients having increased levels of this signature showing poorer disease outcomes (Fig. 4J and Fig. S4I).

Taken together, our results identify Pdk1^high^ colorectal cancer cells as a subpopulation of quiescent cells endowed with stem cell/tumor initiating potential and characterized by a pro-inflammatory and mesenchymal phenotype involved in tumor progression.

### Pdk1^high^ cells drive chemoresistance in CRC

Our results indicate that Pdk1^high^ cells exhibit several features associated to chemoresistance and tumor relapse, including increased glycolysis, a slow-cycling/quiescent phenotype, tumor-initiating potential and activation of inflammation/IFN-related pathways (*37–40*). Based on this, we hypothesized that Pdk1-dependent metabolic heterogeneity could be a key factor contributing to CRC chemoresistance. To test this possibility, we first treated MC38-*Pdk1*-mCherry spheroids with 5-Fluorouracil (5-FU), a frontline therapy for CRC patients, and analyzed cell death in Pdk1^high^ and Pdk1^low^ cells. Whereas more than 35% of Pdk1^low^ cells were dead after 3 days of treatment, 5-FU did not induce cell death in Pdk1^high^ cells in these conditions (Fig. 5A). Concomitantly, 5-FU treatment resulted in an expansion of Pdk1^high^ cells (Fig. 5B).

**Figure 5.**
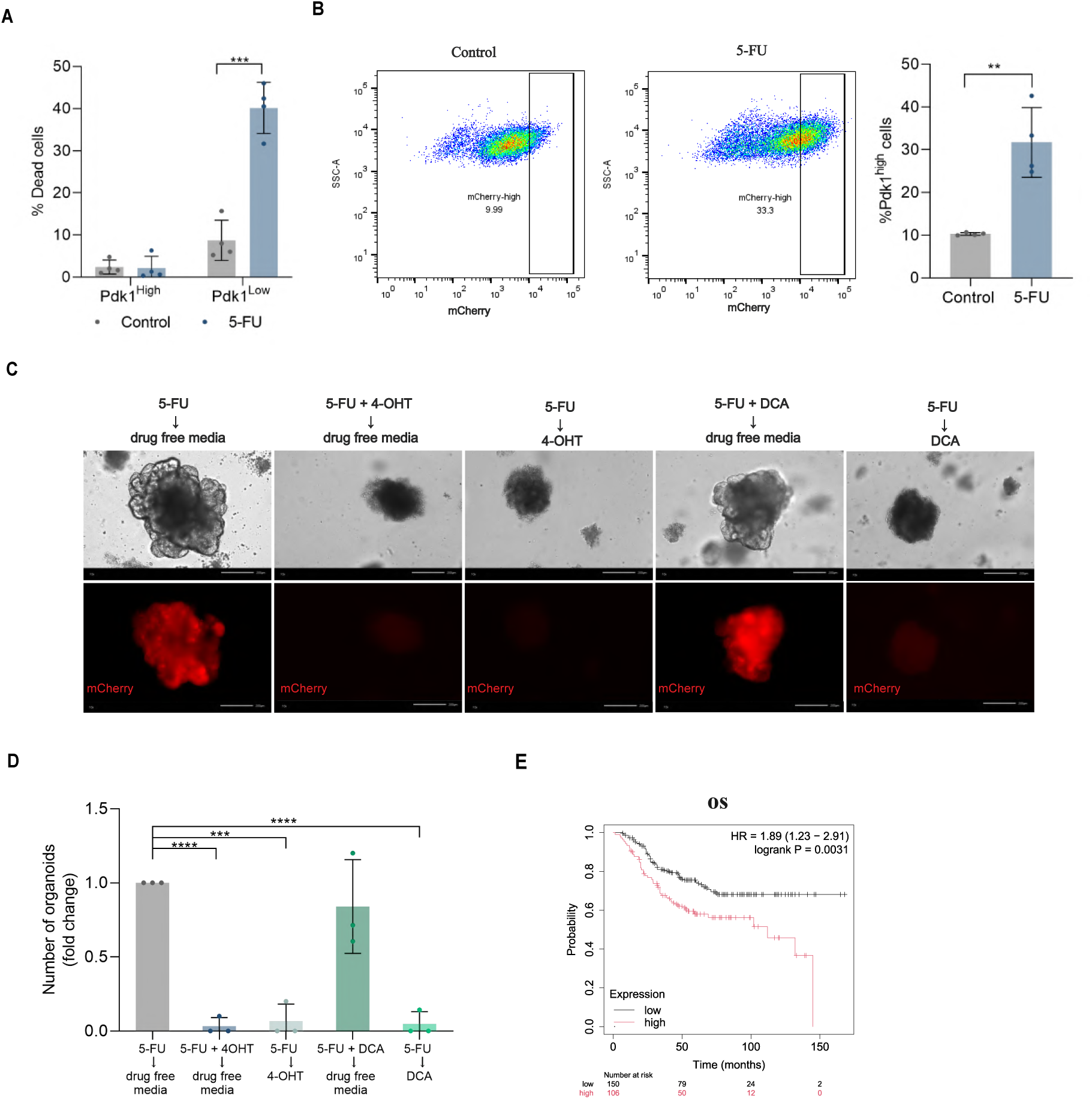
Pdk1^high^ cells drive chemoresistance in CRC. **(A)** Quantification of the percentage of dead cells (DAPI-positive) in Pdk1^high^ and Pdk1^low^ populations FACS-sorted from MC38-Pdk1-mCherry spheroids treated with vehicle (control) or 5-FU (50 µM) for 72 hours (n = 4). Data are expressed as mean ± SEM. Statistical significance was assessed using unpaired t-test, ***p < 0.001. **(B)** Representative flow cytometry plots showing the shift in mCherry fluorescence intensity (mCherry signal versus SSC-A) in MC38-Pdk1-mCherry spheroids treated with vehicle (control) or 5-FU (50 µM) for 72 hours, illustrating the relative enrichment of the Pdk1^high^ population following treatment, together with quantification of the percentage of mCherry^high^ cells in control and 5-FU-treated conditions across 4 independent experiments (n = 4). **(C)** Representative brightfield and mCherry fluorescence images of organoids at experimental endpoint for each of the five treatment conditions, illustrating the differential recovery capacity across groups. Scale bar, 200μm. **(D)** Quantification of the number of recovered organoids, expressed as fold change relative to the 5-FU → drug-free condition, in PDOs subjected to 5-FU treatment followed by the indicated recovery conditions (n = 3). Data are expressed as mean ± SEM. Statistical significance was assessed using unpaired t-test, ***p < 0.001, **** p < 0.0001. **(E)** Kaplan-Meier analysis of overall survival (OS) in CRC patients after adjuvant chemotherapy stratified by high or low expression of the Pdk1^high^ gene signature.

To further explore the contribution of chemoresistant Pdk1^high^ cells to tumor relapse, we treated PDOs-*PDK1*-mCh/Cre/DTA with 200μM of 5-FU during 15 days, a condition that induces the death of most of the cells within the organoids (fig. S5A), and analyzed organoid formation in 5-FU-free media. After around 30 days of 5-FU washout, we observed the emergence of incipient organoids containing mostly mCherry^high^ cells (fig. S5B), which were able to develop into fully mature organoids (Fig. 5C), suggesting that surviving PDK1^high^ cells could be responsible for organoid regrowth after chemotherapy. To test this, we treated PDOs with 4-OHT (to ablate PDK1^high^ cells) during 5-FU treatment and analyzed organoid regrowth in drug-free media. Strikingly, organoid formation was severely impaired in these conditions (Fig. 5C and 5D), indicating that PDK1^high^ cells are required for organoid formation after 5-FU therapy. Similarly, ablation of surviving PDK1^high^ cells during the recovery phase (after 5-FU treatment) almost completely abrogated organoid formation (Fig. 5C and 5D), highlighting the prominent role of these cells during tumor recurrence. Lastly, we assessed whether the metabolic properties of PDK1^high^ cells could be involved in their chemoresistant phenotype. To do so, we performed organoid formation experiments as above in the presence of DCA. Interestingly, we found that DCA inhibited organoid regrowth only when administered in the recovery phase (Fig. 5C and 5D), suggesting that PDK activity is required for organoid formation but not for 5-FU chemoresistance. This result is in line with our findings showing impaired stem cell/tumor initiating potential upon PDK1 inhibition (Fig. 3I).

Kaplan-Meir analysis of patient survival after adjuvant chemotherapy showed decreased overall survival (OS) in those patients expressing high levels of the Pdk1^high^ signature (Fig. 5E), suggesting poorer responses to chemotherapy. Indeed, both relapse-free survival (RFS) and post-progression survival (PPS) were negatively correlated with the expression of the Pdk1^high^ signature (fig. S5C and S5D), indicating a higher probability of tumor relapse and the appearance of more aggressive tumors in CRC patients with high levels of expression of this gene signature.

Taken together, our results indicate that Pdk1^high^ cells represent a chemoresistant cell subset within colorectal tumors involved in tumor relapse.

### Mitochondrial remodeling links the Pdk1^high^ state to the activation of IFN signaling

Having established the phenotype of Pdk1^high^ cells and the contribution of these cells to CRC progression, we sought to provide mechanistic insights on how the metabolic state of Pdk1^high^ cells could determine their phenotype. According to the RNA-seq data from MC38 tumors, activation of type I and type II IFN pathways and downregulation of mitochondrial metabolism are the most prominent phenotypic features of Pdk1^high^ cells. Indeed, we found decreased expression of several genes involved in mitochondrial metabolism and function (*Slc25a5, Tomm40, Ndufa4*) in Pdk1^high^ sorted cells from MC38-*Pdk1*-mCherry spheroids, whereas key ISGs (*Ifit3, Ifit1, Stat1, Ifih1*) were upregulated in these cells (Fig. 6A). These results confirm the expression data obtained in our RNA-seq experiment and, importantly, indicate that the phenotypic features defining Pdk1^high^ cells are cell-intrinsic, suggesting an intimate link between their metabolic and cellular phenotypes.

**Figure 6.**
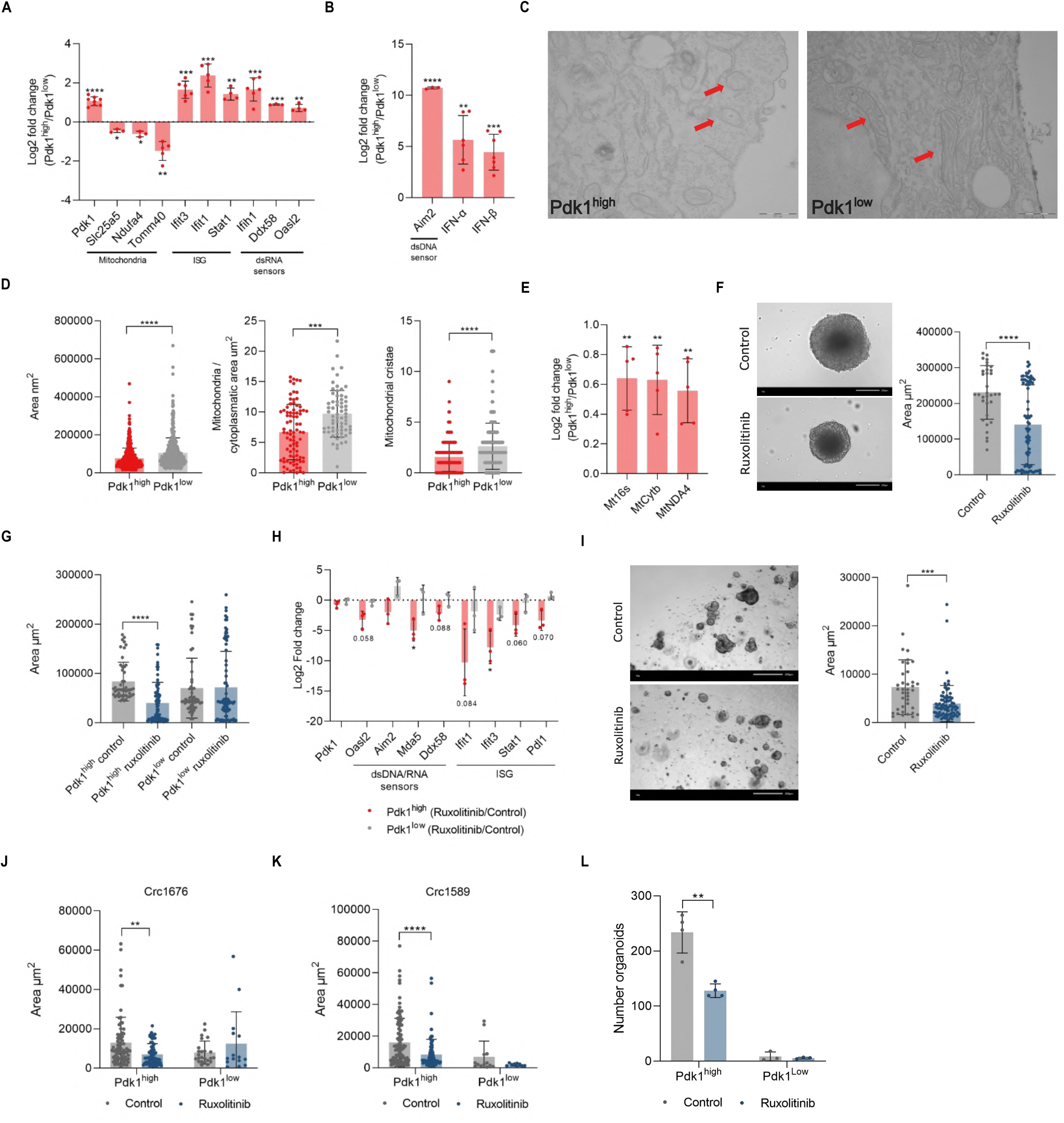
Mitochondrial remodeling links the Pdk1^high^ state with the activation of IFN signaling. **(A)** RT-qPCR analysis of mitochondrial genes, ISG and dsRNA sensors in Pdk1^high^ and Pdk1^low^ cells FACS-sorted from MC38-Pdk1-mCherry spheroids. Data are expressed as log₂ fold change (Pdk1^high^/Pdk1^low^) (n = 3–8). Data are expressed as mean± SEM. Statistical significance was assessed using a one-sample t-test, *p < 0.05, **p < 0.01, ***p<0.001. **(B)** RT-qPCR analysis of *Aim2* and type I IFNs in Pdk1^high^ and Pdk1^low^ cells FACS-sorted from MC38-Pdk1-mCherry spheroids. Data are expressed as log₂ fold change (Pdk1^high^/Pdk1^low^) (n = 3–7). Data are expressed as mean± SEM. Statistical significance was assessed using a one-sample t-test, *p < 0.05, **p < 0.01, ***p<0.001, ****p<0.0001. **(C)** Representative TEM images of Pdk1^high^ cells and Pdk1^low^ cells sorted from MC38-Pdk1-mCherry spheroids and replated for 24 hours in 2D culture. Arrows indicate mitochondria. Scale bar, 500 nm. **(D)** Quantification of mitochondrial morphological parameters in Pdk1^high^ and Pdk1^low^ cells, including mitochondrial area (nm²), number of mitochondria per cell area, and number of cristae per mitochondrion (n = 530 and n = 430 mitochondria analyzed in Pdk1^high^ and Pdk1^low^ cells, respectively). Data are expressed as the mean ± SD. Statistical significance was assessed using a Mann-Whitney test, ***p < 0.001, ****p < 0.0001. **(E)** RT-qPCR quantification of mitochondrial DNA species in cytosolic fractions of Pdk1^high^ and Pdk1^low^ cells FACS-sorted from MC38-Pdk1-mCherry spheroids. DNA levels of mitochondrial-encoded genes Mt16s, mt-Cytb, and mt-Nd4 were measured as markers of cytosolic mitochondrial DNA accumulation. Data are expressed as log₂ fold change (Pdk1^high^/Pdk1^low^) (n = 5). Data are expressed as mean± SEM. Statistical significance was assessed using a one-sample t-test, **p < 0.01. **(F)** Representative brightfield images and quantification of spheroid area (µm²) in MC38-Pdk1-mCherry spheroids treated with vehicle (control) or ruxolitinib (5 µM) at 96 hours. Each data point represents individual spheroids from 4 independent experiments. Data are expressed as mean± SEM. Statistical significance was assessed using unpaired t-test, ****p < 0.0001. **(G)** Quantification of spheroid area (µm²) of spheroids generated exclusively from FACS-sorted Pdk1^high^ or Pdk1^low^ cells and treated with vehicle (control) or ruxolitinib (5 µM). Each data point represents individual spheroids from 3 independent experiments. Data are expressed as mean± SD. Statistical significance was assessed using unpaired t-test, ****p < 0.0001. **(H)** RT-qPCR analysis of innate immune and interferon-stimulated genes expressed as log₂ fold change comparing ruxolitinib-treated versus vehicle-treated conditions in Pdk1^high^ cells-derived and Pdk1^low^ cells-derived spheroids (n = 3). Data are expressed as mean ± SEM. Statistical significance was assessed using a one-sample t-test, *p < 0.05. **(I)** Representative brightfield images and quantification of organoid area (µm²) of CRC1589 PDOs treated with vehicle (control) or ruxolitinib (5 µM) from single-cell suspensions (n = 3). Data are expressed as mean ± SD. Statistical significance was assessed using unpaired t-test, ***p < 0.001. **(J)** Quantification of organoid area (µm²) of CRC1676 PDOs generated from FACS-sorted Pdk1^high^ or Pdk1^low^ cells and treated with vehicle (control) or ruxolitinib (5 µM) (n = 3). Data are expressed as mean ± SD. Statistical significance was assessed using unpaired t-test, **p < 0.01. **(K)** Quantification of organoid area (µm²) in CRC-1589 generated from FACS-sorted Pdk1high or Pdk1low cells and treated with vehicle (control) or ruxolitinib (5 µM) (n = 3). Data are expressed as mean ± SD. Statistical significance was assessed using unpaired t-test, ****p < 0.0001. **(L)** Quantification of organoid number in CRC1676 and CRC1589 PDOs treated with vehicle (control) or ruxolitinib (5 µM) from single-cell suspensions (n = 3). Data are expressed as mean ± SEM. Statistical significance was assessed using unpaired t-test, **p < 0.01.

Cell-intrinsic activation of IFN pathways has been associated with the presence of cytosolic nucleic acids, including dsDNA and dsRNA, which are sensed by pattern recognition receptors (PRR) from the Toll-like receptor (TLR), NOD-like receptor (NLR) and retinoic acid-inducible gene I (RIG-I)-like receptor (RLR) families, as well as multiple intracellular DNA sensors (*41*). Of note, among the upregulated ISGs in Pdk1^high^ cells, we found the dsRNA sensors *Ifih1* (MDA5), *Ddx58* (RIG-1) and *Oasl2* (Fig. 6A, fig S4E and fig. S6A). Notably, one of the most upregulated genes in Pdk1^high^ cells from MC38 tumors was the dsDNA sensor *Aim2* (fig. S4E), which we confirmed to be highly expressed in Pdk1^high^ cells from spheroids (Fig. 6B). Moreover, cleaved caspase-1, the proteolytically activated form of caspase-1 and the hallmark effector of Aim2, was detected in Pdk1^high^ cells and was almost absent in Pdk1^low^ cells (fig. S6A), providing direct biochemical evidence for Aim2 inflammasome activation. Since mitochondrial stress has been shown to trigger the activation of type I IFN responses through the release of mitochondrial DNA (mtDNA) to the cytosol (*42*), we speculated that high activity of Pdk1 could link mitochondrial stress to the accumulation of cytosolic mtDNA in Pdk1^high^ cells, thus leading to the activation of a type I IFN response in these cells. Indeed, we found increased expression of *<u>Ifna</u>* and *Ifnb* in Pdk1^high^ cells sorted from MC38 spheroids (Fig. 6B). To test the mechanistic link between high Pdk1 levels and IFN signaling, we first analyzed mitochondrial structure in Pdk1^high^- and Pdk1^low^-sorted cells from MC38-*Pdk1*-mCherry spheroids. To minimize the morphological changes associated with sorting-induced cellular stress, we established 2D cultures from mCherry sorted cells from spheroids. Analysis of pPDH and Pdk1 protein levels 24 hours after plating revealed that sorted Pdk1^high^ cells maintained elevated Pdk1 and pPDH levels compared to Pdk1^low^ cells, comparable to the levels observed in cells sorted directly from spheroids (fig. S6B), while this difference was lost at longer time points (fig. S6C). Of note, analysis of PDH activity both just after cell sorting and after 24h of 2D culture confirmed the metabolic status of Pdk1^high^ cells, that is, low PDH activity (fig. S6D). Pdk1^high^ and Pdk1^low^ cells were then subjected to transmission electron microscopy (TEM) to directly examine mitochondrial ultrastructure at single-cell resolution. Our results showed decreased numbers of mitochondria in Pdk1^high^ cells, which were also smaller in size and, importantly, contained less cristae (Fig. 6C and 6D). Consistently, we observed decreased protein levels of Timm44 and Opa1 in Pdk1^high^ cells (fig. S6E). These structural differences are in line with the transcriptomic evidence of altered mitochondrial function in Pdk1^high^ cells and suggested that downregulation of mitochondrial OxPhos and structural gene expression might be accompanied by morphological alterations of mitochondria, changes that are consistent with mitochondrial stress, and that may underlie the release of mitochondrial dsDNA into the cytosol. Indeed, cell fractionation experiments showed increased levels of cytosolic mtDNA in Pdk1^high^ cells (Fig. 6E and fig. S6F), thus providing a mechanistic explanation for the activation of the innate immune and interferon transcriptional programs that define this cell population.

Altogether, these data indicate that mitochondrial remodeling in Pdk1^high^ cells leads to the release of mtDNA to the cytosol triggering the activation of IFN signaling, and supports a role for Pdk1-dependent cellular metabolism as a regulator of sterile inflammation in cancer cells.

### Blocking IFN signaling inhibits the tumorigenic properties of Pdk1^high^ cancer cells

To functionally validate the activity of the IFN signaling pathway in Pdk1^high^ cells, we treated MC38-Pdk1-mCherry spheroids with ruxolitinib, a selective inhibitor of JAK1 and JAK2 that blocks downstream STAT1 and STAT2 phosphorylation and abrogates transcriptional responses to type I and type II IFNs. Treatment with ruxolitinib resulted in an approximately 40% reduction in spheroid size compared to controls (Fig. 6F). Notably, ruxolitinib treatment had a significantly greater impact on spheroids derived from Pdk1^high^ cells compared to those derived from Pdk1^low^ cells (Fig. 6G), as evidenced by a more pronounced reduction in spheroid size, indicating that activation of IFN signaling in Pdk1^high^ cells contributes to spheroid growth. In line with this, gene expression analysis of Pdk1^high^ cells-derived spheroids treated with ruxolitinib revealed a marked downregulation of ISGs, while the same genes showed minimal or no significant response to ruxolitinib treatment in Pdk1^low^ cells-derived spheroids (Fig. 6H). Altogether, these results demonstrate that the IFN and innate immune signaling program is selectively and functionally active in Pdk1^high^ cells, that it is mediated through the JAK-STAT pathway, and that it contributes to the role of this population in driving the growth of colorectal cancer spheroids.

To extend the functional validation of interferon signaling to a human CRC context, we performed similar experiments with PDOs in the presence or absence of ruxolitinib. In unsorted single-cell suspensions from CRC1676 and CRC1589 organoids, ruxolitinib treatment resulted in a significant reduction in organoid area compared to controls (Fig. 6I). To determine whether this effect was preferentially mediated through PDK1^high^ cells, we performed organoid formation assays from FACS-sorted PDK1^high^ and PDK1^low^ populations from these PDO lines. Ruxolitinib treatment caused a significant reduction in the size of organoids derived from PDK1^high^ cells, while no changes were observed in PDK1^low^ cells-derived organoids (Fig. 6J-K), indicating that the IFN pathway is selectively active and functionally relevant in PDK1^high^ cells in PDOs. Of note, ruxolitinib treatment impaired the organoid formation capacity of PDK1^high^ cells (Fig. 6L). In contrast, PDK1^low^ cells, which already display an intrinsically reduced capacity to form organoids compared to PDK1^high^ cells, showed no significant difference in organoid number (Fig. 6L) upon ruxolitinib treatment. These results are consistent with an active role of the IFN pathway in driving tumor-initiating potential of PDK1^high^ cells.

Taken together, these results validate in a human CRC model the preferential dependence of PDK1^high^ cells on JAK-STAT signaling, and further support the existence of an active and functionally important IFN transcriptional program specifically within this tumor-initiating population.

## Discussion

Besides genetic heterogeneity, non-genetic mechanisms have been described to drive ITH. Here, we describe the presence of metabolically diverse cancer cell populations within colorectal tumors, which establish an intratumoral metabolic heterogeneity in CRC based on the expression levels of the glycolytic enzyme Pdk1. Our phenotypic and functional characterization of Pdk1^high^ cells revealed that high levels of Pdk1 determines a cell state characterized by quiescence, stem cell-like potential, EMT and activation of IFN pathways, which regulate tumor progression, resistance to chemotherapies and tumor relapse.

Tumor metabolic heterogeneity has been mostly attributed to differential nutrient and oxygen availability to cancer cells (*12*). In glioblastoma, cancer cells residing in perivascular regions exhibit increased oxidative phosphorylation and anabolic metabolism compared to hypoxic regions (*43*), while the expression of phosphoglycerate dehydrogenase in breast cancer cells inversely correlates with tumor vascularization (*8*). This metabolic heterogeneity between normoxic and hypoxic areas can be recapitulated in 3D spheroids, where TCA cycle metabolites accumulate in outer regions while glycolytic intermediates exhibit higher levels in the hypoxic core (*44*). However, our LSR-3D imaging and spatial analyses showed no differences in the distribution of Pdk1^high^ and Pdk1^low^ cancer cells relative to blood vessels. In line with this, we found that Pdk1^high^ cells are distributed throughout spheroids and patient-derived organoids, with no selective accumulation at the core where nutrients and oxygen are scarce. Although we cannot completely rule out that Pdk1 expression might be increased in some cells due to hypoxia, our results suggest that, overall, the Pdk1^high^ state represents a cell-intrinsic metabolic program that can be adopted independently of the local oxygen and nutrient microenvironment. Indeed, histopathological analysis of orthotopic colorectal tumors revealed that PDK1^high^ cells are not uniformly distributed but are instead enriched at the invasive front and, especially, in PDCs, specific anatomical regions strongly associated with lymph node metastasis, lymphovascular invasion, and poor prognosis in CRC (*17*). Intriguingly, this spatial enrichment in PDCs is accompanied by an increase in the proportion of Ki67^+^ PDK1^high^ cells, suggesting a quiescence to proliferation switch as cells transition from the tumor core to the invasive periphery. We observed a similar heterogeneous pattern of PDK1 expression in liver metastatic lesions, indicating that newly formed tumors in this tissue maintain PDK1-driven metabolic heterogeneity. Also in this case, the proliferative status of PDK1^high^ cells changes dynamically as metastatic lesions grow. In micrometastases, approximately 10% of PDK1^high^ cells are Ki67^+^, suggesting that these cells actively contribute to the early stages of metastatic colonization by re-entering the cell cycle in the new metastatic niche. However, it could also reflect the higher percentage of PDK1^high^ cells in PDCs, which can be released to the circulation and seed the liver. In this context, PDK1 has been described to drive metastasis formation in the liver by breast and colorectal cancer cells (*45, 46*), further supporting a potential role of PDK1^high^ cells in the establishment of CRC metastasis. As metastatic lesions increase in size, however, the proportion of Ki67^+^ PDK1^high^ cells progressively decreases, consistent with a return to quiescence as a new tumor hierarchy is re-established. The ability to transit between quiescent and proliferative states, rather than being fixed in either, indicates that PDK1^high^ cells are not in an irreversible cell cycle arrest state and that their quiescent phenotype might be reversible, which can be compatible with the stem cell-like and tumor relapse properties of these cells. Indeed, we found increased expression of *Hes1*, a known regulator of quiescence reversibility (*47*), in Pdk1^high^ cells (fig.S4E).

Beyond our spatial analyses, transcriptomic and functional experiments further support a role for Pdk1^high^ cells in driving tumor progression and aggressiveness. The enrichment of Pdk1^high^ cells within PDCs is consistent with their transcriptional identity, which display prominent upregulation of *Zeb1*, *Zeb2*, and other EMT drivers, and have been associated to tumor budding and PDCs (*48*). Moreover, the Pdk1^high^ signature from both mouse tumors and human PDOs is associated to decreased overall survival of CRC patients, and our genetic ablation experiments confirm that Pdk1^high^ cells contribute to tumor growth. Importantly, our organoid formation assays indicate that these cells drive CRC organoid formation, which, together with our previous results in mouse models (*13*), is in agreement with a stem cell-like and tumor initiating potential of PDK1^high^ cells, suggesting that they could contribute not only to tumor growth but also to tumor initiation. In line with this, a recent report has linked high glycolysis to a cancer stem cell (CSC) phenotype in CRC. Machine learning integration of bulk and single-cell RNA-seq data has identified a high glycolytic state (HGS) associated to higher CytoTRACE scores, consistent with HGS cells having stem cell-like properties (*49*). However, our results suggest that Pdk1^high^ cells are molecularly distinct from other previously described cancer stem cell identities, as they do not express LGR5, fetal-like or regenerative stem cell transcriptional programs. This is consistent with the growing recognition that the CSC compartment in CRC is not homogeneous, but comprises multiple, molecularly distinct subpopulations that coexist within the same tumor and can interconvert in response to microenvironmental and therapeutic cues (*50*). The identification of Pdk1^high^ cells as a functionally important and molecularly unique CSC population therefore adds an important new dimension to our understanding of intratumoral heterogeneity in colorectal cancer.

A key finding of our study is the requirement of Pdk1^high^ cells for organoid re-growth after cytotoxic chemotherapy, suggesting a role for these cells as drivers of tumor relapse in CRC. Several lines of evidence suggest that Pdk1^high^ cells might represent a pre-existing chemoresistant population within colorectal tumors. First, they are present in primary tumors, even at very early stages. Second, they display well-known phenotypic features associated to a chemoresistant state, such as a slow-cycling/quiescence state, tumor-initiating potential and enhanced glycolysis (*37, 38, 40*). Similarly, a recent work has described upregulation of transcriptional programs associated to IFN responses and JAK-STAT signaling in metastatic colorectal tumors resistant to 5-FU-based chemotherapies (*51*), further supporting that Pdk1^high^ cells exhibit properties of a chemoresistant state. Lastly, our results indicate that Pdk1^high^ cells show intrinsic resistance to 5-FU-induced apoptosis and that their genetic ablation, concurrently with chemotherapy or during the recovery phase, or inhibition of Pdk1 activity with DCA during the recovery phase, impairs organoid formation, suggesting a direct role of pre-existing, surviving Pdk1^high^ cells on tumor relapse. Indeed, our results are consistent with the reported ability of DCA to sensitize CRC cells to 5-FU-based chemotherapy, to the positive correlation of PDK4 expression with 5-FU resistance of CRC cells and the lower expression of several subunits of the mitochondrial F1F0-ATP synthase in 5-FU-resistant cells (*52–55*). Conversely, a recent study has shown that 5-FU treatment promotes mitochondrial biogenesis and metabolism in CRC cells, which was associated with increased cell survival (*56*). However, these same cells also showed higher glycolytic activity upon 5-FU treatment, as reflected by the increased incorporation of glucose carbons into pyruvate, lactate and alanine, consistent with reduced flux through PDH. Moreover, although our results indicate that the mitochondrial network is altered in Pdk1^high^ cells, we cannot rule out that other carbon sources could be feeding the TCA cycle to maintain mitochondrial metabolism. Indeed, increased fatty acid oxidation has been associated with both PDH inhibition and mitochondrial stress (*57, 58*) and, interestingly, upregulation of a mitochondrial fatty acid metabolism gene signature has been recently associated with poorly responsive colorectal tumors (*51*).

Lastly, our work provides a mechanistic connection between the metabolic and phenotypic states of Pdk1^high^ cells. Based on our results, we propose that Pdk1-driven metabolic reprogramming leads to mitochondrial stress and the release of mitochondrial nucleic acids to the cytosol, which in turn promotes a cell-intrinsic IFN response and AIM2-dependent inflammasome activation. While AIM2 and type I IFN signaling have been usually associated with antitumor immune responses (*59–61*), emerging evidence indicates that chronic activation of these pathways within cancer cells can paradoxically exert pro-tumoral effects. AIM2 inflammasome activation has been linked to tumor angiogenesis, radioresistance, migration and PD-L1 expression in oral squamous carcinoma (*62*), whereas sustained IFN signaling in cancer cells has been described to promote stemness, EMT properties and resistance to immune checkpoint blockade (*22, 63*). Interestingly, this sustained IFN signaling is associated with the acquisition of an epigenetic inflammatory memory maintained by STAT1 and IRF3, driving the expression of a subset of IFN-stimulated genes (*63*), most of which, including STAT1, are upregulated in Pdk1^high^ cells (Fig.4D and Fig. S4E). Similarly, AIM2 has been described as a regulator of inflammatory memory in skin epithelial stem cells (*64*). Together with our results, these observations suggest a link between sustained inflammatory pathways and the tumorigenic properties of Pdk1^high^ cells. Importantly, our experiments with ruxolitinib provide functional data establishing JAK-STAT/IFN signaling as selective functional dependency of Pdk1^high^ cells in both murine and human CRC models, highlighting the Pdk1-IFN axis as therapeutically actionable target.

Taken together, our results establish Pdk1^high^ cells as key contributors of tumor initiation and progression, therapy resistance and tumor relapse in CRC, and identify metabolic heterogeneity as a functionally important and clinically relevant dimension of intratumoral heterogeneity. Furthermore, we found that the Pdk1^high^ metabolic state is cell intrinsic and determines a defined cellular phenotype characterized by stem cell-like activity, EMT properties and sustained activation of inflammatory pathways, which are associated with mitochondrial stress-induced release of mitochondrial nucleic acids to the cytosol and activation of AIM2 inflammasome and IFN signaling. Genetic ablation of Pdk1^high^ cells, Pdk1 inhibition or JAK-STAT pathway blockade impair the tumorigenic properties of this cell population, thus establishing a clinically relevant connection between Pdk1-driven metabolic heterogeneity and IFN signaling in driving aggressive CRC.

## Material and Methods

### Mice

All mice were maintained at the animal facility of the Candiolo Cancer Institute-FPO and the Parc Científic de Barcelona (PCB). Animal care and experimental protocols were performed in compliance with the Ethical Commission of the Candiolo Cancer Institute, the Italian Ministry of Health (Approval# 185/2018-PR), the Ethics Committee on Animal Experimentation (CEEA) of the PCB (Project#20-046) and the Government of Catalunya (Approval#11238). Five-to-seven weeks old male and female C57BL6/J mice used in this study were obtained from our established colony (founders were purchased from Janvier Laboratories). Five-to-seven weeks old male and female NOD-SCID mice (NOD.CB17-Prkdc^scid^/NCrCrl) were obtained from an established colony at the animal facility of the Candiolo Cancer Institute (founders were purchased from Charles River Laboratories).

### Cell lines and CRC Patient-Derived Organoids

MC38 and HCT116 colorectal cancer cell lines used in this study were kindly provided by Prof. Alberto Bardelli (Candiolo Cancer Institute-FPO). Cell lines were cultured under standard conditions (37°C, 5% CO₂) in DMEM-high glucose (Merck, D5796) supplemented with 10% FBS, 1% P/S, and 2 mM L-Glutamine. Cells were maintained in 10 cm diameter plates and were passaged when they reached approximately 90% confluency, typically every 3–4 days. 3D spheroids were generated from MC38-Pdk1-mCherry/CreERT2-FlexDTA cells or HCT116-PDK1-mCherry/CreERT2-FlexDTA cells using the hanging drop method. Briefly, 40 μl drops containing 2500 cells were deposited onto the inner surface of inverted 15 cm culture dish lids. Plates were carefully inverted and incubated at 37°C with 5% CO₂ for 3–4 days to allow spheroid formation. After the incubation period, spheroids were harvested by tilting the lid and washing gently with PBS to facilitate detachment. PDOs are part of the XENTURION collection (*15*). Four lines were used for in vitro experiments: three derived from primary colorectal tumors (CRC1676, CRC1898 and CRC1707) and one derived from a liver metastasis harboring a KRAS G12C mutation (CRC1589). Organoids were cultured in 50 μl domes of Basement Membrane Extract (BME-Cultrex) (R&D Systems, 3533-005-02,) in 24-well plates, overlaid with 500 μL of PDOs Medium: Basal Crypt Medium (Advanced DMEM/F12) (Gibco, 12634010) supplemented with 10 mM HEPES (Corning, 15323661), 2 mM L-Glutamine, 1% Penicillin/Streptomycin, 1× N2 supplement (Gibco, 17502-048), 1× B27 supplement (Gibco, 17504-044), and 20 ng/mL recombinant EGF (Gibco, PMG8043). Medium was refreshed every 3 days and organoids were passaged every 5–7 days. For organoid formation experiments, PDOs were collected and dissociated to single cells with TrypLE™ Express for 15 minutes at 37°C. 1000 cells were resuspended in 50 μl of BME and plated in 24-well plates in PDOs medium supplemented with ROCK inhibitor for 2–3 days, after which medium was replaced with standard PDOs medium for continued growth.

### Constructs and viral infection

The Pdk1-mCherry reporter was previously generated (*13*). Briefly, a 1.3 kb region upstream of the ATG of the mouse *Pdk1* gene was amplified by PCR and cloned into the pHAGE2.1-EF1aL-mCherry-W vector (kindly provided by Dr. Darrell Kotton) via AgeI/NotI restriction sites, generating the pHAGE2.1-Pdk1-mCherry construct. Two versions of this construct were generated using the same vector backbone: one with the mouse Pdk1 promoter and another one with the human PDK1 promoter. To generate the Pdk1-CreERT2 reporter, the CreERT2 coding sequence was amplified by PCR from the pCAG-CreERT2 plasmid (Addgene, #14797) using primers incorporating 5’-NotI and 3’-BamHI restriction sites. Before cloning, an internal BamHI site within the CreERT2 coding sequence was eliminated by site-directed mutagenesis. The PCR-amplified CreERT2 fragment was then cloned into pHAGE2.1-Pdk1-mCherry by replacing the mCherry sequence via NotI/BamHI restriction-based cloning, yielding the pHAGE2.1-Pdk1-CreERT2 construct. Also in this case, the human version was generated following the same strategy. To generate the pMSCV-flexDTA construct, the flexDTA cassette (loxP–DTA–loxP) was first subcloned from pAAV-mCherry-flexDTA (Addgene, #58536) into pGEM-T-easy (Promega) by TA cloning. The flexDTA cassette was subsequently transferred into the retroviral backbone pMSCV-puro by EcoRI-based restriction cloning. Viral particles containing the above-mentioned plasmids were synthesized using lentiviral (pCMV-dR8.91) packaging plasmid with pCMV-VSV-G (Addgene) and concentrated by ultracentrifugation.

For MC38 and HCT116 cells infection, cells were seeded in 6-well plates and grown to 50–70% confluency. Culture medium was then replaced with fresh medium containing 5–10 μL of concentrated viral particles and 10 μg/mL polybrene (Sigma-Aldrich, TR-1003). Cells were incubated overnight and, the following day, viral medium was replaced with standard culture medium. MC38 cells and HCT116 cells transduced with the Pdk1-mCherry construct were selected by fluorescence-activated cell sorting (FACSAria™ Fusion, BD Biosciences) based on mCherry fluorescence. Cells transduced with the flexDTA construct were selected by puromycin resistance (Sigma-Aldrich, P7130). For experiments involving Pdk1-CreERT2 transduction, cells were always co-transduced with Pdk1-mCherry to enable monitoring of transduction efficiency, assuming comparable infection rates for both constructs. The transduction of PDOs was performed following a modified protocol based on Onuma et al (*65*). PDOs were collected and dissociated to single cells with TrypLE™ Express for 15 minutes at 37°C. 1-5×10^5^ cells were resuspended in 500 μl of PDOs medium supplemented with 20 μM ROCK inhibitor (Y-27632) containing 10 μg/mL polybrene and 5–10 μl of the concentrated virus. The mixture was centrifuged at 600 rpm for 1 hour at 4°C (spinoculation) and subsequently incubated at 37°C with 5% CO₂ for 4 hours. Cells were resuspended in 50 μl of BME and plated in 24-well plates in PDOs medium supplemented with ROCK inhibitor for 2–3 days, after which medium was replaced with standard PDOs medium for continued growth. For PDK1-mCherry-transduced PDOs, individual clones were selected to ensure that mCherry expression levels faithfully reflected endogenous PDK1 promoter activity rather than differences in transduction efficiency. PDOs transduced with the flexDTA construct were selected by puromycin resistance. For experiments involving PDK1-CreERT2 transduction, PDOs were always co-transduced with PDK1-mCherry to enable monitoring of transduction efficiency, assuming comparable infection rates for both constructs.

### Genetic ablation experiments

For genetic ablation experiments, spheroids were treated with 4-hydroxytamoxifen (4-OHT; 1 µM) (Sigma-Aldrich, H6278) from the onset of spheroid formation, allowed to grow for 3–4 days and subsequently used for downstream analyses, including size measurement, immunofluorescence and FACs-sorting analysis. Brightfield and mCherry images of spheroids were acquired using a Nikon Eclipse Ts2R microscope. PDOs were allowed to establish and grow for 3 days, after which the culture medium was supplemented with 4-OHT (1 µM). Organoids were monitored by brightfield and fluorescence microscopy every 24 hours for four consecutive days using a Nikon Eclipse Ts2R fluorescence microscope. At each time point, organoid size and mCherry fluorescence intensity were recorded. When required by the experimental design, organoids were subsequently collected for downstream analyses, including immunofluorescence, FACS analysis, qPCR and protein extraction and western blot. Spheroids and PDOs area was quantified using ImageJ software (NIH). Briefly, the scale was set according to the scale bar present in each image. The contour of each spheroid was then manually traced using the freehand selection tool, and the area was calculated using the built-in “Measure” function of ImageJ.

### Flow cytometry and cell sorting

Flow cytometric analysis and fluorescence-activated cell sorting of Pdk1^high^ and Pdk1^low^ cells from spheroids and PDOs were performed on a FACSAria™ Fusion instrument (BD Biosciences). The gating strategy was applied sequentially as follows (fig. S7). Cells were first identified by SSC-A versus FSC-A. Doublet exclusion was then performed using two sequential gates: FSC-W versus FSC-A, and SSC-W versus SSC-A. Live cells were identified by DAPI exclusion using the V450/50 channel (VIO 450/50-A) versus FSC-A, gating on DAPI-negative events. Within the live single-cells gate, mCherry positive fluorescence was detected using the Green 610/20-A channel. Two populations were defined based on mCherry fluorescence intensity: mCherry^high^ (Pdk1^high^) cells, corresponding to the top 10–15% of mCherry positive events with the highest fluorescence intensity, and mCherry^low^ (Pdk1^low^) cells, corresponding to the bottom 10–15% of events with the lowest mCherry fluorescence. These two populations were sorted and collected for downstream experimental applications.

### 5-FU treatment of spheroids and viability assays

MC38-Pdk1-mCherry spheroids were treated with 5-Fluorouracil (5-FU; 50 µM) (Sigma-Aldrich, F6627) for 3 days from the onset of spheroid formation. Spheroids were collected and dissociated into single-cell suspensions by incubation with TrypLE™ Express (Gibco, 12604021) for 20–25 minutes at 37°C. The cell suspension was then filtered through a 40 µm cell strainer to remove cell aggregates and debris prior to flow cytometric analysis. Cell viability was assessed by flow cytometry on a FACSAria™ Fusion (BD Biosciences) using 4′,6-diamidino-2-fenilindol (DAPI 10 µg/ml) as a live/dead discriminator. Within the DAPI-negative (viable) and DAPI-positive (dead) gates, cells were further stratified based on mCherry fluorescence intensity into Pdk1-mCherry^high^ and Pdk1-mCherry^low^ subpopulations, allowing quantification of the relative survival rates of each population in response to 5-FU treatment. For apoptosis experiments, dissociated cells were resuspended in annexin binding buffer and stained using the Annexin V-FITC Apoptosis Staining/Detection Kit (Abcam, ab14085). Briefly, 2 µL of DAPI (1 µg/mL) and 2 µL of Annexin V-FITC were added to each sample immediately before acquisition. Within each viability gate, cells were further stratified based on mCherry fluorescence intensity into Pdk1-mCherry^high^ and Pdk1-mCherry^low^ subpopulations, enabling quantification of the apoptotic response of each population to 5-FU treatment.

### 5-FU treatment and recovery experiments in PDOs

Three days after routine passaging, organoids were treated with 5-FU (200 µM) for approximately 15 days, until around 98% cell death was achieved. To maintain consistent drug exposure, culture medium was replaced once per week with fresh medium containing 5-FU (200 µM), resulting in two rounds of drug administration over the treatment period. Cell viability was monitored at day 3, day 7, and day 15 by flow cytometric analysis using DAPI as a live/dead discriminator on an Aurora spectral flow cytometer (Cytek). In parallel, organoids were inspected daily by brightfield and fluorescence microscopy to track morphological changes and mCherry fluorescence, the latter serving as a proxy for PDK1^high^ cells viability. To investigate whether concurrent PDK1 targeting modulates chemotherapy response and subsequent recovery, a subset of wells was treated simultaneously from the onset with either 5-FU + 4-OHT (1 µM) or 5-FU + DCA (10 mM). Once around 98% cell death was confirmed, 5-FU was withdrawn and wells were transitioned to one of the following drug-free or treatment conditions depending on their prior treatment history: wells previously treated with 5-FU alone were switched to either drug-free medium, medium supplemented with 4-OHT (1 µM), or medium supplemented with DCA (10 mM); wells previously treated with 5-FU + 4-OHT or 5-FU + DCA were switched to drug-free medium. Following drug withdrawal, organoids were monitored daily by brightfield and fluorescence microscopy over approximately 35–40 days to detect the reappearance of viable, mCherry-positive structures, indicative of organoid relapse. At the experimental endpoint, DAPI-based viability analysis was repeated on the Aurora spectral flow cytometer to quantify the proportion of surviving and recovering cells across all conditions.

### Ruxolitinib treatment and spheroids/organoids formation assays

MC38-Pdk1-mCherry cells were used to generate spheroids in the presence or absence of ruxolitinib (5 µM) (Biotechne, 7064), a pharmacological inhibitor of JAK1/2. Ruxolitinib was dissolved in DMSO and added to the culture medium at the onset of spheroid formation using the hanging drop method. After 3– 4 days, spheroids were collected and imaged, and spheroid area was quantified using ImageJ software (NIH). In some experiments, spheroids were first generated and dissociated into single-cell suspensions, and mCherry^high^ and mCherry^low^ cells were isolated by FACS. Sorted populations were then used to generate new spheroids independently, under control conditions or in the presence of ruxolitinib (5 µM). PDOs were dissociated into single-cell suspensions and 1000 cells per well were plated in BME under standard PDO medium (control) or PDO medium supplemented with ruxolitinib (5 µM). In some experiments, mCherry^high^ and mCherry^low^ populations were first isolated by FACS and 1000 cells from each sorted population were independently plated in BME under control or ruxolitinib-treated conditions. Organoid number and area were monitored at days 3, 7, and 15 post-plating and quantified using ImageJ software (NIH).

### Mouse *in vivo* experiments

Subcutaneous tumors were generated by injecting 3×10^5^ MC38-Pdk1-mCherry/CreERT2-flexDTA or HCT116-PDK1-mCherry/CreERT2-flexDTA cells into the flanks of C57BL6/J or NOD-SCID mice, respectively. For *in vivo* ablation experiments, mice received intraperitoneal injections of tamoxifen (2 mg per injection) every two days beginning three days after tumor cell inoculation and continuing until experimental endpoint, when tumors were excised, weighed, and processed for histological analysis. Orthotopic colorectal tumors were established by intercaecal injection of MC38-Pdk1-mCherry-CreERT2-FlexDTA or HCT116-PDK1-mCherry/CreERT2-flexDTA cells into C57BL6/J or NOD-SCID mice. Animals were maintained under isoflurane anesthesia throughout the procedure. Analgesia was provided by subcutaneous administration of meloxicam (5 mg/kg; Metacam®) at the nape of the neck and lidocaine (2 mg/kg) locally at the abdominal incision site. A midline laparotomy was performed to externalize the caecum, and 5×10^5^ cells, resuspended in 50 μl of PBS, were injected into the cecal tip, with care taken to avoid leakage into the lumen. The abdomen was closed by suture and animal welfare was monitored for five consecutive days post-surgery, with continued meloxicam administration every 12h for 3 days. Caeca were collected approximately one month post-injection for downstream analysis.

### Large Scale Single-cell Resolution 3D imaging

Subcutaneous and orthotopic tumors were collected and processed following a protocol adapted from Rios et al.(*14*). Briefly, tissues were fixed in 4% PFA for 45 minutes at room temperature. After fixation, the tissues were washed for 10 minutes in PBS containing 0.1% Tween-20 (PBST) and incubated in Wash Buffer (WB: PBS, 0.1% Tween-20, 50 μg/mL Ascorbic Acid, 0.05 ng/mL L-Glutathione Reduced) for 2 hours, followed by a 2-hour incubation in Wash Buffer 1 (WB1: PBS, 0.2% Tween-20, 0.2% Triton, 0.02% SDS, 0.2% BSA, 50 μg/mL Ascorbic Acid, 0.05 ng/mL L-Glutathione Reduced) on a roller-mixer at 4°C. Anti-CD31-AlexaFluor 647 (BioLegend, 102515) diluted in WB2 (PBS, 0.1% Triton, 0.02% SDS, 0.2% BSA, 50 μg/mL Ascorbic Acid, 0.05 ng/mL L-Glutathione Reduced) was used to detect blood vessels. The antibody was added after the incubation with WB1 for a minimum of 24 hours at 4°C in a vertical orbital platform. After antibody incubation, tissues were washed twice with WB2 for 1 hour each. Finally, tissues were cleared for optimal imaging by immersing them in FunGI clearing solution (50% Glycerol (v/v), 2.5 M Urea, 2.5 M Fructose, 10.6 mM Tris Base, 1 mM EDTA) and stored at 4°C until further analysis. Confocal imaging was performed on a Carl Zeiss LSM 900 confocal microscope (Zeiss).

### Multiplex-IF and spatial analyses

Opal multiplex immunofluorescent staining on the Leica Bond RX platform was performed by the IRB-Barcelona Histopathology service on formalin-fixed paraffin-embedded tissue sections (3-4 µm in thickness) using the following antibodies and Opal reagents (Akoya, NEL871001KT): anti-RFP/mCherry (Rockland, 600-401-379; ER2, 20’, 1:500) – Opal 570; anti-Ki67 (Abcam, ab15580; ER2, 20’, 1:500) – Opal 620; anti-EpCAM (E6V8Y) (Cell Signaling, 93790; ER2, 20’, 1:500) – Opal 690; and anti-CD31 (Abcam, ab28364; ER2, 20’1:200) – Opal 480. Immunofluorescence images were acquired with a Phenoimager HT digital scanner (Akoya Biosciences) with a 20X objective. DAPI signal of the sample has been acquired with an exposure time of 8.52 ms, and exposure times of the remaining biomarkers were the following: 8.03 ms for CD31-Opal 480; 22.97 ms for RFP-Opal 570; 27.07 ms for Ki67-Opal 620; 110.52 ms for EpCAM-Opal 690. Spatial analysis in different tumor areas and liver metastases was performed using QuPath (*66*). Within annotated areas (ROIs), cells were segmented based on DAPI using the StarDist extension for fluorescent cell detection, which generated fluorescence intensities for each channel on the single cell level. For calculating the % of Pdk1^high^ cells in tumor cores, invasive fronts and poorly differentiated clusters (PDCs) per mouse, single cell fluorescent mCherry signals of all ROIs were pooled, sorted, and the threshold for cells within the 7th percentile of highest mCherry expression across all tumor areas was determined. With this threshold, the % of Pdk1^high^ expressing cells was determined for each area individually. For cell classification analyses, Pdk1^high^, Pdk1^low^ and Ki67+ cells were classified by creating single measurement classifiers, and double positive cells were defined as those passing both single classifiers, identified by applying the classifiers sequentially. An initial intensity threshold was determined for each channel in a representative mouse; the percentile corresponding to this threshold was then calculated and applied uniformly to classify cells in all subsequent mice. In tumor cores, Pdk1^high^ cells were defined as those with mCherry intensity above the 93rd percentile (top 7 % of expressors); Pdk1^low^ were defined as those below the 7th percentile; Ki67+ cells were defined as those above the 70th percentile. In invasive fronts and PDCs, Pdk1^high^ cells were defined as those with mCherry intensities above the 67th percentile; Pdk1^low^ were defined as those below the 33rd percentile; Ki67+ cells were defined as those above the 46th percentile. For comparing metastases of different sizes, the following percentiles were applied: above the 93rd percentile for Pdk1^high^ cells; below the 7th percentile Pdk1^low^ cells; above the 70th percentile for Ki67+ cells; and above the 77th percentile EpCAM+ cells. To identify positive cells for each marker combination (Pdk1^high^ or Pdk1^low^ / Ki67+ or Ki67 - / EpCAM+ or EpCAM-), we developed a program on R 4.5.1 using “quantile” function in stats (v3.6.2) package for percentile determination. QuPath “detection measurements” were exported and used as input, after StarDist cell segmentation. For distance measurements between Pdk1^high^ and Pdk1^low^ cells and CD31+ blood vessels, the same thresholds were applied to classify Pdk1^high^ and Pdk1^low^ cells (above the 93rd percentile and below the 7th percentile, respectively). CD31+ blood vessels were classified using the pixel thresholder. Distances were measured applying “Distances to annotations 2D” in QuPath.

### Immunofluorescence

Spheroids were collected in 15 mL conical tubes, washed once with 1× PBS, and fixed in 4% paraformaldehyde (PFA) for 30 minutes at room temperature (RT). Spheroids were then permeabilized in 1× PBS containing 1% Triton X-100 for 20 minutes at 4°C and blocked with Blocking Solution (1× PBS, 1% BSA, 3% goat serum, 0.2% Triton X-100) for 1 hour at 4°C. Spheroids were incubated with primary antibodies diluted 1:100 in Antibody Solution (1× PBS, 0.1% BSA, 0.3% goat serum, 0.2% Triton X-100) overnight at 4°C. After two 5-minute washes in 1× PBS, secondary antibodies (1:500 in Antibody Solution) were applied for 1 hour at RT. Nuclei were counterstained with DAPI (10 µg/ml) for 30 minutes at RT. Following two final PBS washes, spheroids were mounted on glass slides with 50 μl of FluoroMount-G™ and stored protected from light at 4°C until imaging. PDOs were stained following the same procedure after collection from BME by incubation with Cell Recovery Solution (Corning, 354253) for 30 minutes at 4°C. Tissue immunofluorescence experiments were done as previously described (*13*). Briefly, rehydrated slides were washed and subjected to antigen retrieval in citrate buffer pH 6 at 95°C for 20 min, blocked with TSA blocking reagent (Perkin Elmer FP1020) and incubated with primary antibodies in blocking reagent overnight at 4°C. After three washes with TNT buffer (100mM Tris-HCl pH 7.5, 150mM NaCl, 0.05% Tween-20), slides were incubated with secondary antibodies in blocking buffer for 1h at room temperature and counterstained with DAPI. Primary antibodies used include: anti-phospho-PDH-E1a-Ser293 (AbCam, ab92696), anti-mCherry (Invitrogen, m11217), anti-Ki67 (Invitrogen, 14-5698-82; Cell Signaling, 9449). AlexaFluor-488/555/647 conjugated secondary antibodies were purchased from Invitrogen. Confocal imaging of spheroids, PDOs, and tissue sections was performed on a Carl Zeiss LSM 900 confocal microscope (Zeiss) at the core facilities of the University of Barcelona (CCiT-UB).

### Protein extraction and Western blot

Protein extracts were prepared from sorted cell pellets resuspended in 200 μL of RIPA lysis buffer (50 mM Tris-HCl pH 7.5, 0.5% Deoxycholic acid, 0.1% SDS, 150mM NaCl, 5mM EDTA) supplemented with protease and phosphatase inhibitors. Samples were sonicated on ice (2 cycles of 5 minutes) and centrifuged at 10000 × g for 20 minutes at 4°C. Protein samples were resolved on pre-cast 4–20% Bis-Tris gradient gels (Millipore, MP42G15) in 1 X MES-SDS running buffer (Thermo Fisher Scientific, J62138.K2) and transferred onto nitrocellulose membranes using iBlot 2 NC Mini or Regular stacks (Invitrogen, IB23001/2) using the iBlot 2 dry blotting system (Invitrogen). Membranes were blocked for 1 hour at RT in TBS-T [1× TBS, 0.1% Tween-20] containing 5% Bovine Serum Albumin (BSA) (Thermo Fisher Scientific, BP9701) and incubated overnight at 4°C with primary antibodies diluted 1:1000 in blocking solution. After three washes with TBS-T (5 minutes each), membranes were incubated with the appropriate HRP-conjugated secondary antibodies (1:5000 in blocking solution) for 1 hour at RT. Signal detection was performed using ECL Select Western Blotting Substrate (Thermo Fischer Scientific, 32209) and imaged on an Odyssey Fc imaging system (LI-COR). Primary antibodies used include: anti-Caspase 1 (Novus Biologicals, NB100-56565), anti-Mda5 (Proteinteck, 217151ap), phospho-PDH-E1a-Ser293 (AbCam, ab92696), anti-Pdk1 (Cell Signaling, C47H1), anti-Timm44 (Proteinteck, 133591ap), anti-Tom20 (Selleck Chemicals, F3252), anti-Opa1 (BD Transduction Laboratories, AB_399888), anti-Histone H3 (AbCam, ab1791), anti-Gapdh (Selleck Chemicals, F0003), anti-Vinculin (Santa Cruz, sc73614).

### RNA extraction and Real-Time PCR

Total RNA was extracted from organoids using the RNeasy Mini Kit (Qiagen, 74104) according to the manufacturer’s instructions. Reverse transcription was performed on 1 µg of RNA using the QuantiTect® Reverse Transcription Kit (Qiagen, 205311) according to the manufacturer’s instructions. RT-qPCR was performed on cDNA diluted 1:4 in RNase-free water, using 1× SYBR Green Master Mix (Applied Biosystems, A25742) on a QuantStudio 7 Pro instrument (Thermo Fisher Scientific). Data were analysed using Design & Analysis Software v2 (DA2, Thermo Fisher Scientific). Primers were used at a final concentration of 300 nM). Gene expression was quantified using the relative quantification method. Individual standard curves with known cDNA dilutions were generated for each primer pair, and expression values were interpolated from these curves. Results were normalized to the geometric mean of three housekeeping genes: *Ppib*, *Tbp* and *β-actin*.

### RNA-sequencing: sample preparation

MC38-Pdk1-mCherry subcutaneous tumors (n= 3) were harvested, minced to 1-3mm pieces and digested in 10 ml of digestion buffer (DMEM+0.5mg/ml collagenase B) for 30 minutes at 37°C with rotation. The samples were filtered through a 70 μm cell strainer into a 50 ml tube containing 20 ml of DMEM and centrifuged 5 minutes at 1500 rpm and 4°C. Cell pellets were resuspended in 10 ml of PBS and filtered through a 40 μm cell strainer to get single cells. After centrifugation (1500rpm, 5minutes, 4°C), cells were resuspended in 100 ml of PBS +2% FCS and stained with a cocktail of antibodies containing anti-CD45/CD31/Ter119-FITC (Biolegend) for 30 min on ice. Cells were sorted using a FACSAria™ Fusion based on their mCherry fluorescence. mCherry^high^ and mCherry^low^ populations were defined as the top and bottom 10–15% of CD45/CD31/Ter119 negative cells based on mCherry fluorescence intensity, respectively. For PDOs RNA-seq experiments, three PDO lines harboring the PDK1-mCherry reporter (CRC1676, CRC1898, and CRC1703) were used. For each line, approximately 150 wells of PDOs were grown. 48 hours prior to sorting, ROCK inhibitor (Y-27632; 20 µM) was added to the culture medium to improve single-cell viability during dissociation. On the day of sorting, PDOs were dissociated into single-cell suspensions and sorted on a FACSAria™ Fusion using the gating strategy described above. mCherry^high^ and mCherry^low^ populations were defined as the top and bottom 10–15% of cells based on mCherry fluorescence intensity, respectively. Immediately following sorting, RNA was extracted and samples were submitted to the “Centro Nacional de Análisis Genómico (CNAG)” for library preparation and bulk RNA sequencing.

### RNA-sequencing: data processing and analysis

For samples obtained from MC38-Pdk1-mCherry tumors, RNA-seq reads were mapped with STAR/2.7.8a (*67*) against Mus musculus reference genome (GRCm39). Genes were quantified with RSEM/1.3.0 (*68*) using gencode.vM31 annotation. Differential expression analysis was performed with the limma-voom R package (*69*). Heatmap with the differentially expressed genes (DEGs) was done with the pheatmap R package using the log2cpm values and scaling by row. DEGs were defined as genes with an absolute fold change > 1.5 and an adjusted p-value < 0.1. Multidimensional scaling plot was performed with the plotMDS function. Gene Set Enrichment Analysis (GSEA) of DEGs was performed using GSEA online tool (Broad Institute, Cambridge, MA, USA) to identify enriched biological pathways and gene ontology terms (*70*). Gene Ontology Biological Process (GO: BP) and Hallmark gene set collections from the 2024 release of the Molecular Signatures Database (MSigDB v2024.1) were used (*70, 71*). For samples obtained from PDOs, RNA-seq reads were aligned to the Homo sapiens reference genome (GRCh38) using STAR v2.7.10a with ENCODE-recommended settings. Gene-level quantification was obtained with RSEM v1.3.1 using GENCODE v46 gene annotations. Raw counts were normalized using the trimmed mean of M-values (TMM) method and transformed to log2-counts per million (logCPM). Only genes with at least 1 count per million (CPM) in at least 3 samples were retained for downstream analyses. Differential gene expression analysis was performed with the limma R package v3.54.2 (*72*). The voom function was used to model the mean–variance relationship and compute observation-level precision weights, and these voom-transformed values were used to fit linear models. DEGs were defined as genes with an absolute fold change > 1.5 and an adjusted p-value < 0.1. Multidimensional scaling plots were generated with the limma function plotMDS based on the top 500 genes with the largest absolute log2-fold changes for each pairwise comparison of samples. GSEA of DEGs was performed using GSEA online tool (Broad Institute, Cambridge, MA, USA) to identify enriched biological pathways and gene ontology terms. Gene Ontology Biological Process (GO: BP) and Hallmark gene set collections from the 2024 release of the Molecular Signatures Database (MSigDB v2024.1) were used. Pathway enrichment analyses were done by calculating the gene signature z-score as the sum of z-score values for individual genes contained in the gene signature and expressed in the DEGs list of our RNA-seq divided by the square root of the number of genes.

### Kaplan-Meier survival analysis

To analyze the predictive value of Pdk1^high^ signatures on CRC patient survival, we used the KMplot online tool (https://kmplot.com/analysis/index.php?p=home) (*73*), which contains survival data from 1336 patients from 16 datasets, including overall survival (OS), relapse-free survival (RFS) and post-progression survival (PPS). We used the “multiple genes” option to analyze the correlation of survival data with the expression of all the genes contained in the Pdk1high signature.

### Pyruvate dehydrogenase activity assay

Pyruvate Dehydrogenase (PDH) activity was evaluated using a commercial colorimetric kit (Sigma-Aldrich, MAK567). Cells were collected immediately after sorting or plated into 6-well plates and harvested after 24 hours. Samples were lysed using PDH assay buffer. PDH activity was measured following the manufacturer’s protocol and normalized for the cell number in each well.

### Transmission electron microscopy (TEM)

FACS-sorted mCherryhigh and mCherrylow cells were processed for TEM 24 hours post-sorting following replating in 6-well plates. Fixation was performed by addition of 3 ml of fixative solution (2.5% glutaraldehyde in 1.0 M PHEM buffer) to each sample, followed by two incubations of 30 minutes at room temperature on an orbital shaker at 100 rpm. Cells were then detached using Trypsin-EDTA, transferred to 15 ml conical tubes, and centrifuged at 2000 rpm for 10 minutes. The resulting pellets were washed four times in 0.1 M PHEM buffer, with each wash consisting of resuspension, incubation on an orbital shaker at 100 rpm for 10 minutes at 4°C, and centrifugation at 2000 rpm for 10 minutes. Fixed cell pellets were subsequently submitted to the Electron Cryomicroscopy Unit of the core facilities of the University of Barcelona (CCiT-UB) for TEM sample preparation and imaging. Image analysis, data processing, and quantification were performed using ImageJ/Fiji software (NIH).

### Cytosolic fractionation

Cytosolic fractions were isolated from FACS-sorted Pdk1-mCherry^high^ and Pdk1-mCherry^low^ populations derived from MC38-Pdk1-mCherry spheroids. Sorted cells were centrifuged at 500 × g, washed with PBS, and transferred to 1.5 ml tubes. A fraction of the cells was retained as total homogenate for WB and genomic DNA (gDNA) reference controls. The remaining cells were resuspended in 500 µl of digitonin buffer [150 mM NaCl, 50 mM HEPES pH 7.4, 50 µg/ml digitonin (Sigma-Aldrich, D141)] and incubated for 10 minutes at 4°C to allow selective permeabilization of the plasma membrane while leaving mitochondrial, nuclear, and endoplasmic reticulum membranes intact. Samples were centrifuged at 980 × g for 4 minutes at 4°C, and the supernatant was carefully transferred to a new tube. Samples were centrifuged at 980 x g for 4 minutes at 4°C and the supernatant was carefully transferred to a new tube. The remaining pellet, containing nuclei, mitochondria, and other membrane-bound organelles, was retained as the pellet fraction and processed in parallel for WB and gDNA analysis to serve as a fractionation quality control. A second centrifugation of the supernatant at 980 × g for 4 minutes at 4°C was performed to remove residual debris, followed by a final high-speed centrifugation at 17000 × g for 10 minutes at 4°C to pellet any remaining organellar contamination. The resulting supernatant, representing the cytosolic fraction free of nuclear, mitochondrial, and endoplasmic reticulum content, was collected and divided into aliquots for subsequent qPCR-based detection of mitochondrial dsDNA and WB analysis.

## Acknowledgements

We would like to thank the IRB-Barcelona Histopathology Facility for sample processing and performing Multiplex-IF experiments and the CCiT-UB facilities (Advanced Optical Microscopy, Flow Cytometry, Genomics and Electron Cryomicroscopy) and the Animal Facility of the PCB-UB for technical assistance. We would also like to thank Jorge Lloberas and all the past and current members of the Sebastián lab for their insightful comments and helpful discussions throughout the project.

## Author contributions statement

JC, LP and CS. conceptualized and designed the study; JC. and LP. performed and analyzed most of the experiments; JC., AL, FS, GP and FV performed spheroids and PDOs experiments; LP performed PDOs experiments and QuPath-based spatial analyses, AA-M performed QuPath-based spatial analyses, JC, LP, AM, MS and CS performed orthotopic transplantation experiments; LF, AM, JC and CS performed subcutaneous tumors experiments; AM, JC, LP, AL, GP, FV, FS and ND performed immunostaining on tissues, PDOs and spheroids; EC, AE-C and MG performed and analyzed RNA-seq experiments; CS and AB-D analyzed RNA-seq experiments and performed GSEA analyses; AS provided FFPE human samples and performed histopathological analyses; LT supervised the generation and characterization of the XENTURION PDO collection; ML and DS designed and assisted in mitochondrial studies; JC, LP and CS wrote the manuscript; CS supervised the project.

## References and notes

1. E. Morgan et al., Global burden of colorectal cancer in 2020 and 2040: incidence and mortality estimates from GLOBOCAN. Gut 72, 338–344 (2023).

2. D. A. Lawson, K. Kessenbrock, R. T. Davis, N. Pervolarakis, Z. Werb, Tumour heterogeneity and metastasis at single-cell resolution. Nat Cell Biol 20, 1349–1360 (2018).

3. N. Saoudi González et al., Unravelling the Complexity of Colorectal Cancer: Heterogeneity, Clonal Evolution, and Clinical Implications. Cancers (Basel) 15, (2023).

4. A. Pérez-González, K. Bévant, C. Blanpain, Cancer cell plasticity during tumor progression, metastasis and response to therapy. Nat Cancer 4, 1063–1082 (2023).

5. N. N. Pavlova, J. Zhu, C. B. Thompson, The hallmarks of cancer metabolism: Still emerging. Cell Metab 34, 355–377 (2022).

6. A. Tasdogan et al., Metabolic heterogeneity confers differences in melanoma metastatic potential. Nature 577, 115–120 (2020).

7. J. E. Choi et al., A unique subset of glycolytic tumour-propagating cells drives squamous cell carcinoma. Nat Metab 3, 182–195 (2021).

8. M. Rossi et al., PHGDH heterogeneity potentiates cancer cell dissemination and metastasis. Nature 605, 747–753 (2022).

9. C. Sun et al., Spatially resolved multi-omics highlights cell-specific metabolic remodeling and interactions in gastric cancer. Nat Commun 14, 2692 (2023).

10. Y. Mizuno et al., Intratumoral heterogeneity of glutaminase and lactate dehydrogenase A protein expression in colorectal cancer. Oncol Lett 19, 2934–2942 (2020).

11. Y. Jia et al., Metabolic Heterogeneity of Tumor Cells and its Impact on Colon Cancer Metastasis: Insights from Single-Cell and Bulk Transcriptome Analyses. J Cancer 15, 4175–4196 (2024).

12. M. Demicco, X. Z. Liu, K. Leithner, S. M. Fendt, Metabolic heterogeneity in cancer. Nat Metab 6, 18–38 (2024).

13. C. Sebastian et al., A non-dividing cell population with high pyruvate dehydrogenase kinase activity regulates metabolic heterogeneity and tumorigenesis in the intestine. Nat Commun 13, 1503 (2022).

14. A. C. Rios et al., Intraclonal Plasticity in Mammary Tumors Revealed through Large-Scale Single-Cell Resolution 3D Imaging. Cancer Cell 35, 953 (2019).

15. S. M. Leto et al., XENTURION is a population-level multidimensional resource of xenografts and tumoroids from metastatic colorectal cancer patients. Nat Commun 15, 7495 (2024).

16. J. W. Kim, I. Tchernyshyov, G. L. Semenza, C. V. Dang, HIF-1-mediated expression of pyruvate dehydrogenase kinase: a metabolic switch required for cellular adaptation to hypoxia. Cell Metab 3, 177–185 (2006).

17. T. S. Haddad et al., Tumor budding and poorly differentiated clusters as a biological continuum in colorectal cancer invasion and prognosis. Sci Rep 15, 16944 (2025).

18. Z. Wu, A. E. Autry, J. F. Bergan, M. Watabe-Uchida, C. G. Dulac, Galanin neurons in the medial preoptic area govern parental behaviour. Nature 509, 325–330 (2014).

19. P. Nayak, V. Bentivoglio, M. Varani, A. Signore, Three-Dimensional In Vitro Tumor Spheroid Models for Evaluation of Anticancer Therapy: Recent Updates. Cancers (Basel) 15, (2023).

20. A. S. Qadir et al., CD95/Fas Increases Stemness in Cancer Cells by Inducing a STAT1-Dependent Type I Interferon Response. Cell Rep 18, 2373–2386 (2017).

21. S. Li et al., Interferon alpha-inducible protein 27 promotes epithelial-mesenchymal transition and induces ovarian tumorigenicity and stemness. J Surg Res 193, 255–264 (2015).

22. P. Holicek et al., Type I interferon and cancer. Immunol Rev 321, 115–127 (2024).

23. A. Sakthianandeswaren et al., PHLDA1 expression marks the putative epithelial stem cells and contributes to intestinal tumorigenesis. Cancer Res 71, 3709–3719 (2011).

24. C. Dravis et al., Sox10 Regulates Stem/Progenitor and Mesenchymal Cell States in Mammary Epithelial Cells. Cell Rep 12, 2035–2048 (2015).

25. X. Liang et al., An Enhancer-Driven Stem Cell-Like Program Mediated by SOX9 Blocks Intestinal Differentiation in Colorectal Cancer. Gastroenterology 162, 209–222 (2022).

26. M. Shimokawa et al., Visualization and targeting of LGR5. Nature 545, 187–192 (2017).

27. F. de Sousa e Melo et al., A distinct role for Lgr5. Nature 543, 676–680 (2017).

28. D. K. L. Tsang et al., A single cell survey of the microbial impacts on the mouse small intestinal epithelium. Gut Microbes 14, 2108281 (2022).

29. C. Morral et al., Zonation of Ribosomal DNA Transcription Defines a Stem Cell Hierarchy in Colorectal Cancer. Cell Stem Cell 26, 845–861.e812 (2020).

30. K. Ganesh et al., L1CAM defines the regenerative origin of metastasis-initiating cells in colorectal cancer. Nat Cancer 1, 28–45 (2020).

31. A. Cañellas-Socias et al., Metastatic recurrence in colorectal cancer arises from residual EMP1. Nature 611, 603–613 (2022).

32. E. G. Vasquez et al., Dynamic and adaptive cancer stem cell population admixture in colorectal neoplasia. Cell Stem Cell 29, 1213–1228.e1218 (2022).

33. L. Solé et al., p53 wild-type colorectal cancer cells that express a fetal gene signature are associated with metastasis and poor prognosis. Nat Commun 13, 2866 (2022).

34. A. Fumagalli et al., Plasticity of Lgr5-Negative Cancer Cells Drives Metastasis in Colorectal Cancer. Cell Stem Cell 26, 569–578.e567 (2020).

35. R. C. Mustata et al., Identification of Lgr5-independent spheroid-generating progenitors of the mouse fetal intestinal epithelium. Cell Rep 5, 421–432 (2013).

36. K. Pelka et al., Spatially organized multicellular immune hubs in human colorectal cancer. Cell 184, 4734–4752.e4720 (2021).

37. I. Puig et al., TET2 controls chemoresistant slow-cycling cancer cell survival and tumor recurrence. J Clin Invest 128, 3887–3905 (2018).

38. C. Zhong et al., Chemoresistance mechanisms to 5-Fluorouracil and reversal strategies in lung and breast cancer. Sci Rep 15, 6074 (2025).

39. M. Musella et al., Type I IFNs promote cancer cell stemness by triggering the epigenetic regulator KDM1B. Nat Immunol 23, 1379–1392 (2022).

40. J. Peng et al., Altered glycolysis results in drug-resistant in clinical tumor therapy. Oncol Lett 21, 369 (2021).

41. T. Gong, L. Liu, W. Jiang, R. Zhou, DAMP-sensing receptors in sterile inflammation and inflammatory diseases. Nat Rev Immunol 20, 95–112 (2020).

42. A. Bahat, T. MacVicar, T. Langer, Metabolism and Innate Immunity Meet at the Mitochondria. Front Cell Dev Biol 9, 720490 (2021).

43. S. Kumar et al., Intra-Tumoral Metabolic Zonation and Resultant Phenotypic Diversification Are Dictated by Blood Vessel Proximity. Cell Metab 30, 201–211.e206 (2019).

44. L. E. Flint et al., Characterization of an Aggregated Three-Dimensional Cell Culture Model by Multimodal Mass Spectrometry Imaging. Anal Chem 92, 12538–12547 (2020).

45. W. Qin et al., The double inhibition of PDK1 and STAT3-Y705 prevents liver metastasis in colorectal cancer. Sci Rep 9, 12973 (2019).

46. F. Dupuy et al., PDK1-Dependent Metabolic Reprogramming Dictates Metastatic Potential in Breast Cancer. Cell Metab 22, 577–589 (2015).

47. L. Sang, H. A. Coller, J. M. Roberts, Control of the reversibility of cellular quiescence by the transcriptional repressor HES1. Science 321, 1095–1100 (2008).

48. A. Pavlič et al., Tumour budding and poorly differentiated clusters in colon cancer - different manifestations of partial epithelial-mesenchymal transition. J Pathol 258, 278–288 (2022).

49. Y. Du et al., Machine learning integration of bulk and single-cell RNA-seq data reveals glycolytic heterogeneity in colorectal cancer. Med Oncol 42, 458 (2025).

50. E. Gil Vazquez et al., Dynamic and adaptive cancer stem cell population admixture in colorectal neoplasia. Cell Stem Cell 29, 1612 (2022).

51. M. Avolio et al., The Molecular and Functional Landscape of Resistance to FOLFIRI Chemotherapy in Metastatic Colorectal Cancer. Cancer Discov 16, 270–295 (2026).

52. Y. Liang et al., Dichloroacetate restores colorectal cancer chemosensitivity through the p53/miR-149-3p/PDK2-mediated glucose metabolic pathway. Oncogene 39, 469–485 (2020).

53. J. Tong et al., Synergistic antitumor effect of dichloroacetate in combination with 5-fluorouracil in colorectal cancer. J Biomed Biotechnol 2011, 740564 (2011).

54. D. Leclerc et al., Oncogenic role of PDK4 in human colon cancer cells. Br J Cancer 116, 930–936 (2017).

55. Y. K. Shin et al., Down-regulation of mitochondrial F1F0-ATP synthase in human colon cancer cells with induced 5-fluorouracil resistance. Cancer Res 65, 3162–3170 (2005).

56. D. Y. Moss et al., Mitochondrial metabolism determines chemotherapy sensitivity in colorectal cancer. Nat Metab 8, 1751–1771 (2026).

57. K. T. Chambers et al., Chronic inhibition of pyruvate dehydrogenase in heart triggers an adaptive metabolic response. J Biol Chem 286, 11155–11162 (2011).

58. E. L. Seifert, C. Estey, J. Y. Xuan, M. E. Harper, Electron transport chain-dependent and - independent mechanisms of mitochondrial H2O2 emission during long-chain fatty acid oxidation. J Biol Chem 285, 5748–5758 (2010).

59. S. Dihlmann et al., Lack of Absent in Melanoma 2 (AIM2) expression in tumor cells is closely associated with poor survival in colorectal cancer patients. Int J Cancer 135, 2387–2396 (2014).

60. J. E. Wilson et al., Inflammasome-independent role of AIM2 in suppressing colon tumorigenesis via DNA-PK and Akt. Nat Med 21, 906–913 (2015).

61. L. Zitvogel, L. Galluzzi, O. Kepp, M. J. Smyth, G. Kroemer, Type I interferons in anticancer immunity. Nat Rev Immunol 15, 405–414 (2015).

62. H. W. Chiu et al., AIM2 promotes irradiation resistance, migration ability and PD-L1 expression through STAT1/NF-κB activation in oral squamous cell carcinoma. J Transl Med 22, 13 (2024).

63. J. Qiu et al., Cancer cells resistant to immune checkpoint blockade acquire interferon-associated epigenetic memory to sustain T cell dysfunction. Nat Cancer 4, 43–61 (2023).

64. S. Naik et al., Inflammatory memory sensitizes skin epithelial stem cells to tissue damage. Nature 550, 475–480 (2017).

65. K. Onuma et al., Genetic reconstitution of tumorigenesis in primary intestinal cells. Proc Natl Acad Sci U S A 110, 11127–11132 (2013).

66. P. Bankhead et al., QuPath: Open source software for digital pathology image analysis. Sci Rep 7, 16878 (2017).

67. A. Dobin et al., STAR: ultrafast universal RNA-seq aligner. Bioinformatics 29, 15–21 (2013).

68. B. Li, C. N. Dewey, RSEM: accurate transcript quantification from RNA-Seq data with or without a reference genome. BMC Bioinformatics 12, 323 (2011).

69. C. W. Law, Y. Chen, W. Shi, G. K. Smyth, voom: Precision weights unlock linear model analysis tools for RNA-seq read counts. Genome Biol 15, R29 (2014).

70. A. Subramanian et al., Gene set enrichment analysis: a knowledge-based approach for interpreting genome-wide expression profiles. Proc Natl Acad Sci U S A 102, 15545–15550 (2005).

71. A. S. Castanza et al., Extending support for mouse data in the Molecular Signatures Database (MSigDB). Nat Methods 20, 1619–1620 (2023).

72. M. E. Ritchie et al., limma powers differential expression analyses for RNA-sequencing and microarray studies. Nucleic Acids Res 43, e47 (2015).

73. B. Győrffy, Integrated analysis of public datasets for the discovery and validation of survival-associated genes in solid tumors. Innovation (Camb) 5, 100625 (2024).

