## Supplementary Figures for "Pdk1-dependent metabolic heterogeneity promotes aggressive colorectal cancer through cell-intrinsic interferon signaling"

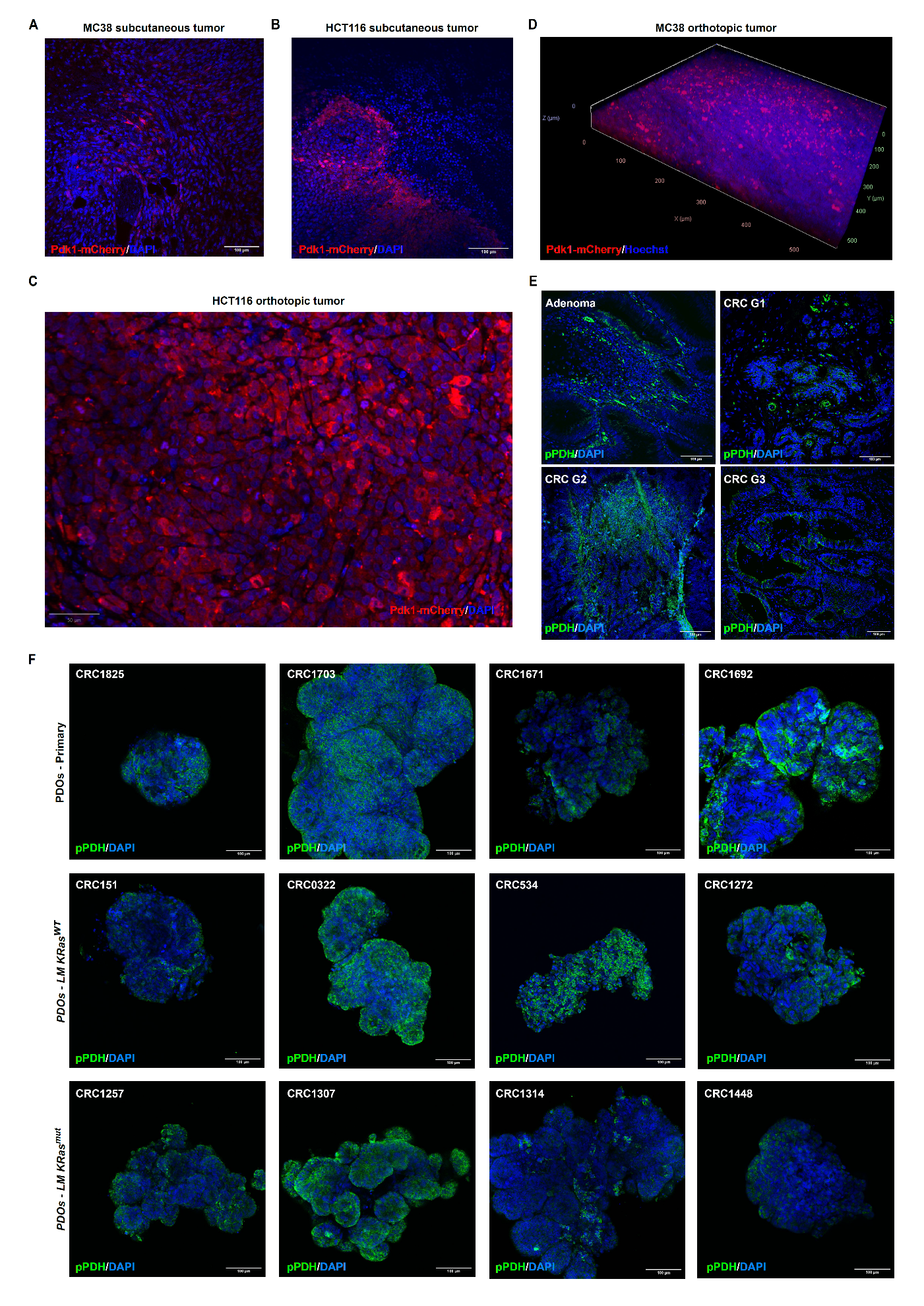

Fig. S1. Pdk1-dependent metabolic heterogeneity in CRC.

(A) Representative fluorescence microscopy image of a subcutaneous colorectal tumor section derived from MC38-Pdk1-mCherry cells. Scale bar, 100μm. (B) Representative fluorescence microscopy image of a subcutaneous colorectal tumor section derived from HCT116-PDK1-mCherry cells. Scale bar, 100μm. (C) Representative fluorescence microscopy image of an orthotopic colorectal tumor section derived from HCT116-PDK1-mCherry cells. Scale bar, 50μm. (D) LSR-3D imaging of an orthotopic tumor derived from MC38-Pdk1-mCherry cells. (E) pPDH staining of tissue sections from human colorectal tumors of different grades. Scale bar, 100μm. (F) pPDH staining of PDOs derived from primary CRC (upper panels), liver metastatic lesions KRasWT (middle panels) and KRasG12C (lower panels). Scale bar, 100μm.

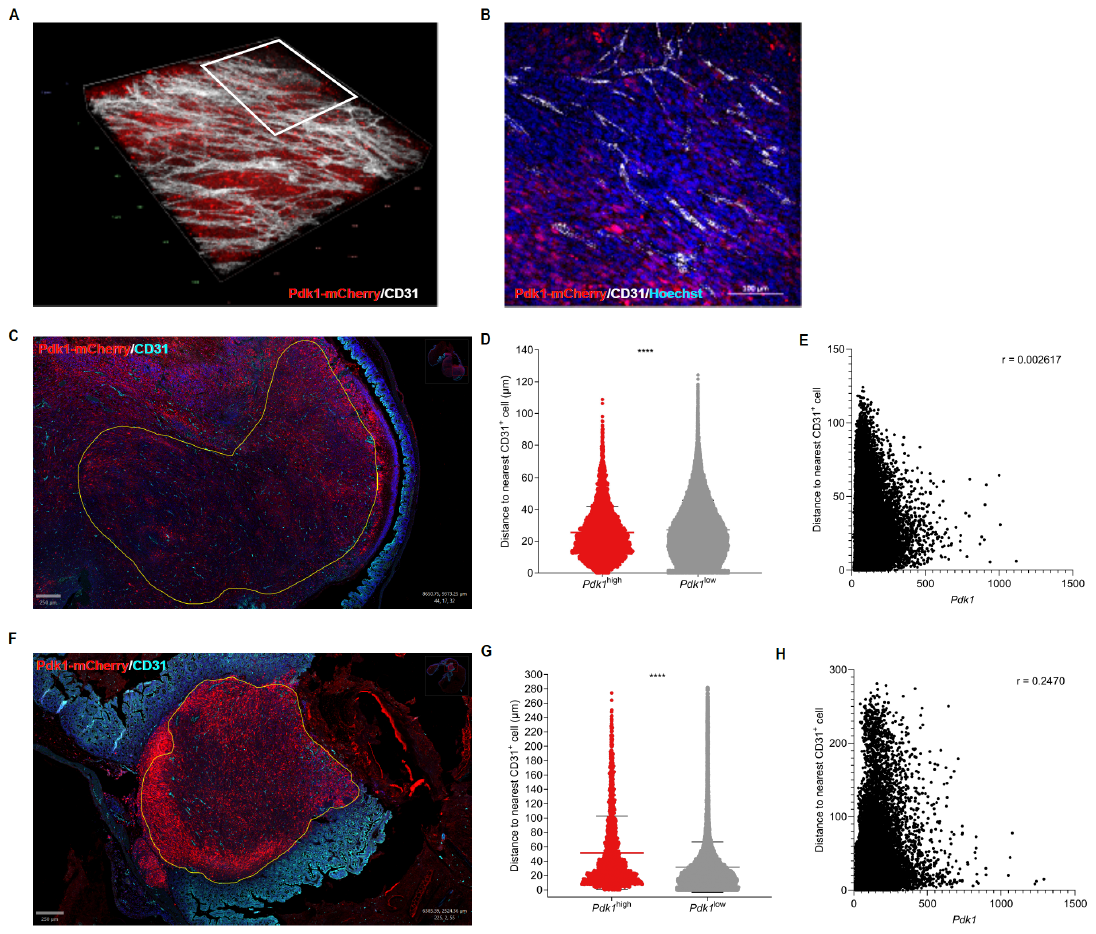

Fig. S2. Spatial distribution of Pdk1^high^ cells relative to blood vessels in colorectal tumors.

(A) LSR-3D imaging of an orthotopic tumor derived from MC38-Pdk1-mCherry cells and stained with anti-CD31 (white). (B) Representative 2D image from a section of the tumor in (A). Scale bar, 100μm. (C) Representative fluorescence microscopy image of an orthotopic MC38-Pdk1-mCherry tumor stained with anti-CD31. Yellow outline indicates the tumor section analyzed in D and E. Scale bar, 250μm. (D) Violin plot showing the distribution of distances (μm) to the nearest CD31^+^ endothelial cell for Pdk1^high^ and Pdk1^low^ cancer cells. (E) Scatter plot showing the correlation between Pdk1 fluorescence intensity (x-axis) and distance to the nearest CD31^+^ cell (y-axis) for individual cancer cells within the tumor. (F) Representative fluorescence microscopy image of a second individual orthotopic MC38-Pdk1-mCherry tumor stained with anti-CD31. Yellow outline indicates the tumor section analyzed in G and H. Scale bar, 250μm. (G) Violin plot showing the distribution of distances (μm) to the nearest CD31^+^ endothelial cell for Pdk1^high^ and Pdk1^low^ cancer cells. (H) Scatter plot showing the correlation between Pdk1 fluorescence intensity (x-axis) and distance to the nearest CD31^+^ cell (y-axis) for individual cancer cells within the tumor.

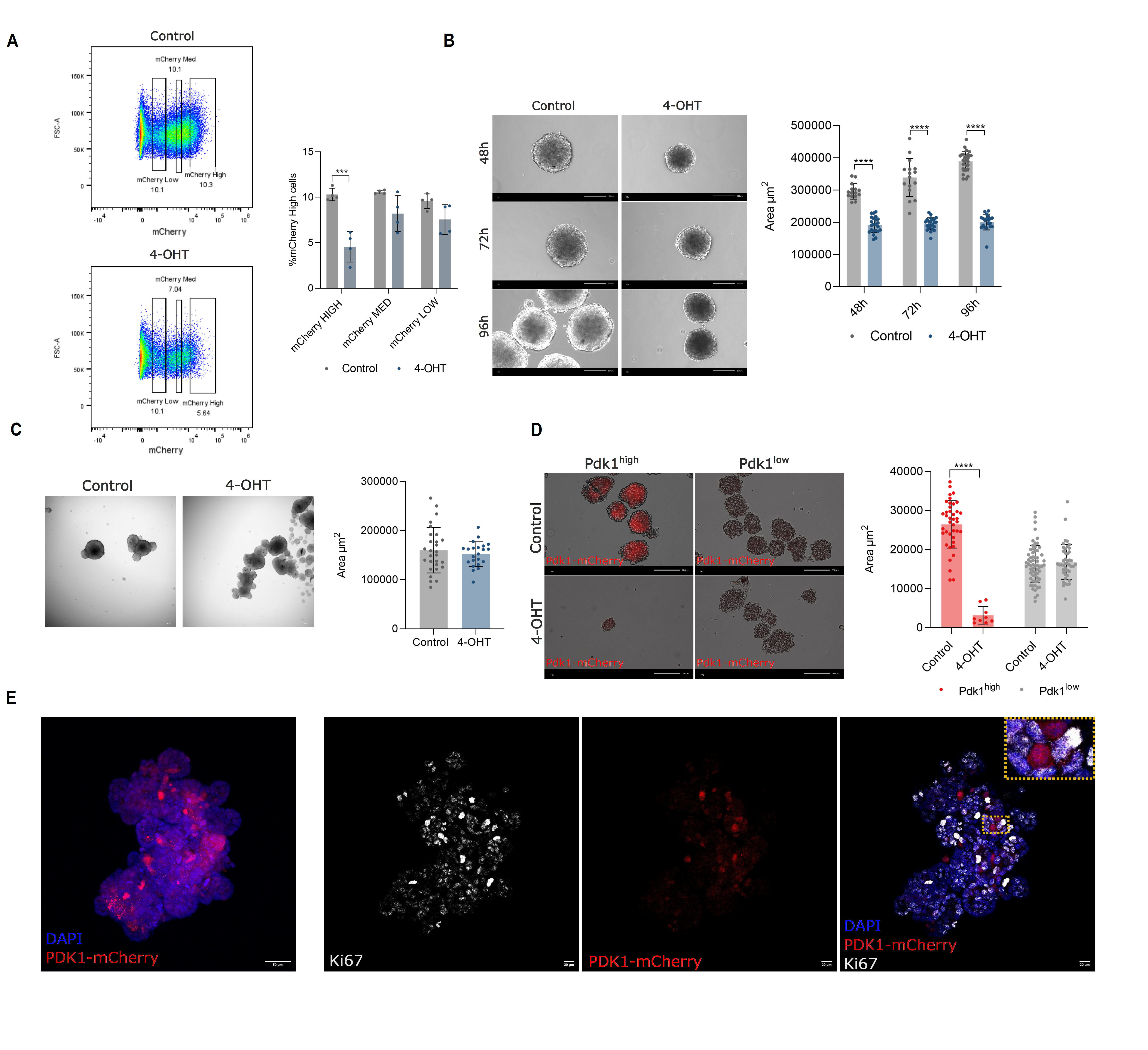

**Fig. S3. The Pdk1-mCherry/CreERT2-flexDTA system selectively eliminates Pdk1^high^ cells.**

(A) FACS analysis of mCherry populations of MC38-Pdk1-mCh/Cre-flexDTA spheroids treated with 4-OHT. Data are expressed as mean± SEM. Statistical significance was assessed using multiple t-test, ***p < 0.001. (B) Representative brightfield images of HCT116-PDK1-mCh/Cre-FlexDTA spheroids treated with vehicle (control) or 4-OHT at 48-, 72-, and 96-hours post-treatment (left panel) and quantification of spheroid area (µm²) (right panel). Data shown are from a representative experiment (n = 3). Data are expressed as mean ± SD. Statistical significance was assessed using multiple t-test, ****p < 0.0001. (C) Brightfield images of MC38-Pdk1-mCh/Cre spheroids (lacking the flexDTA cassette) treated with 4-OHT (left panel) and quantification of spheroid area (right panel) (n=3). Data are expressed as mean± SEM. Statistical significance was assessed using unpaired t-test. (D) Representative brightfield and mCherry fluorescence images of spheroids generated exclusively from FACS-sorted Pdk1-mCherry^high^ or Pdk1-mCherry^low^ cells (right panel) and quantification of spheroid area (µm²) (n= 3). Data are expressed as mean± SD. Statistical significance was assessed using unpaired t-test, ****p < 0.0001. (E) Confocal microscopy images of CRC1589 PDOs co-stained for DAPI, mCherry and Ki67, with a magnified inset highlighting the absence of overlap between individual mCherry-positive cells and Ki67-positive nuclei.

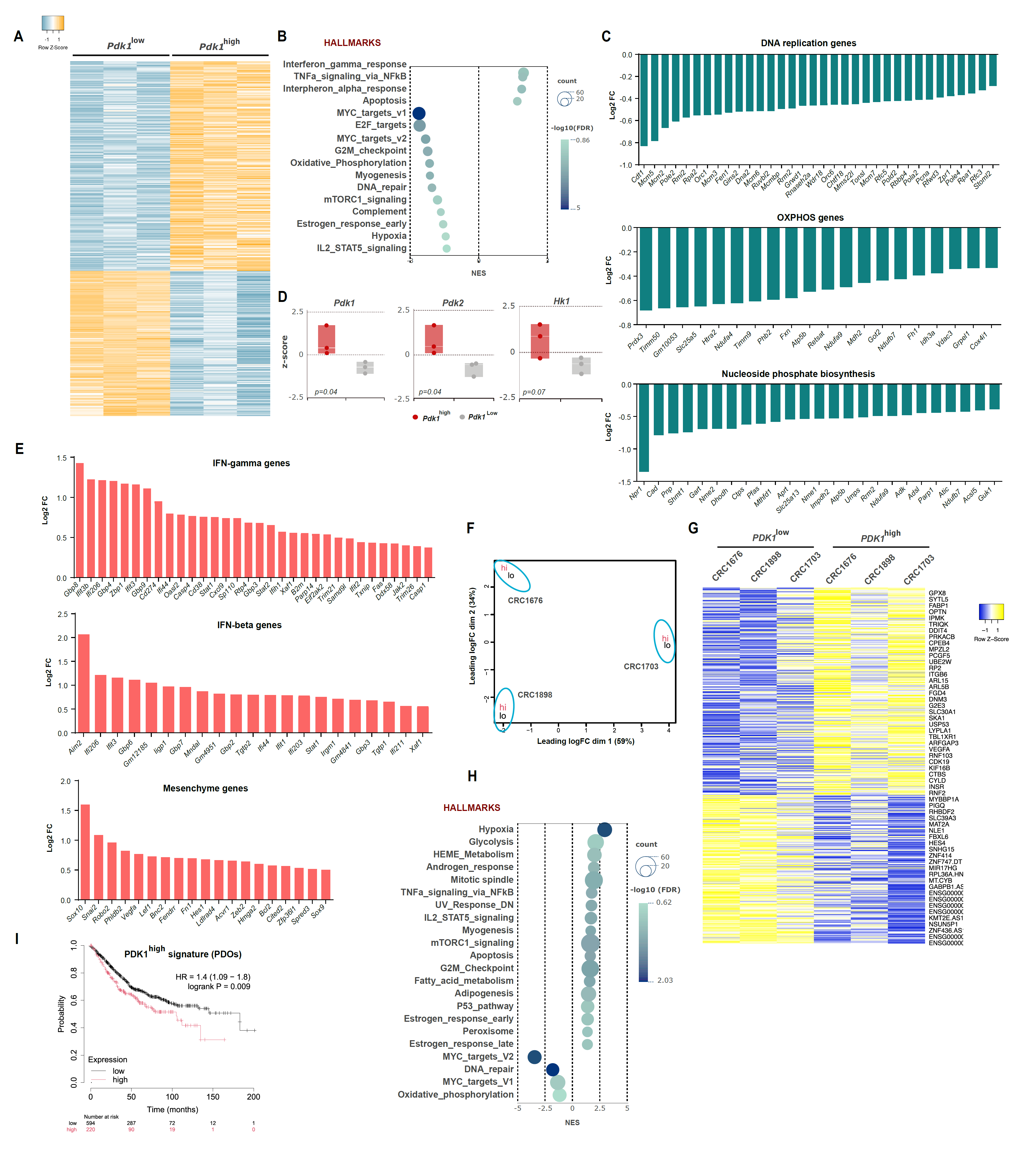

**Fig S4. Transcriptional profiling of Pdk1^high^ cells.**

(A) Heatmap showing the row Z-score of differentially expressed genes between Pdk1^low^ and Pdk1^high^ cells sorted from MC38-Pdk1-mCherry subcutaneous tumors (n=3). (B) Bubble plot showing the normalized enrichment scores (NES) of Hallmarks gene sets significantly upregulated and downregulated in Pdk1^high^ cells compared to Pdk1^low^ cells. Bubble size reflects the number of genes in each gene set and color intensity reflects the -log10(FDR). (C) Bar plots showing the log₂ Fold change (log2FC) of individual genes within three downregulated gene categories in Pdk1^high^ cells: DNA replication genes, OxPhos genes, and nucleoside phosphate biosynthesis genes. (D) Box plots showing the z-score normalized expression levels of key glycolytic genes (*Pdk1, Pdk2* and *Hk1*) in Pdk1^high^ and Pdk1^low^ cells sorted from MC38-Pdk1-mCherry subcutaneous tumors. (E) Bar plots showing the log₂ Fold change (log2FC) of individual genes within upregulated gene categories in Pdk1^high^ cells: IFN-gamma genes, IFN-beta genes and EMT genes. (F) Principal component analysis (PCA) plot of RNA sequencing data from PDK1^high^ and PDK1^low^ cells FACS-sorted from three independent PDO lines (CRC1676, CRC1898, and CRC1703). (G) Heatmap showing the row Z-score of differentially expressed genes between PDK1^high^ and PDK1^low^ cells sorted from the PDO lines in (F). (H) Bubble plot showing the normalized enrichment scores (NES) of Hallmarks gene sets significantly upregulated and downregulated in PDK1^high^ cells compared to PDK1^low^ cells from PDOs. Bubble size reflects the number of genes in each gene set and color intensity reflects the -log10(FDR). (I) Kaplan-Meier survival analysis of overall survival (OS) in colorectal cancer patients stratified by high or low expression of the PDO-PDK1^high^ gene signature (HR = 1.4, 95% CI 1.09–1.8, log-rank p = 0.009). Numbers at risk are shown below the graph.

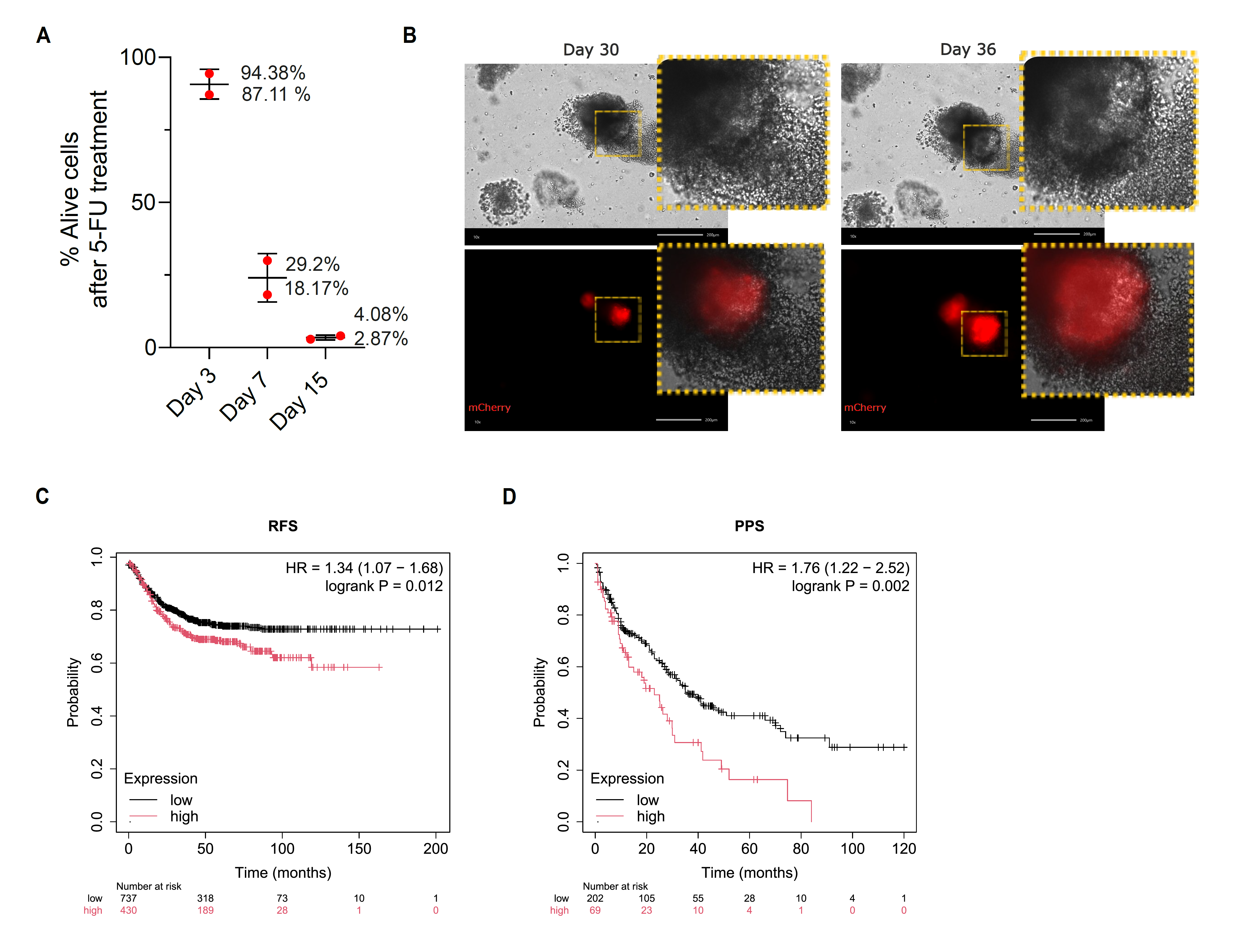

**Fig S5. PDK1^high^ cells are involved in tumor relapse after 5-FU treatment in PDOs.**

(A) Flow cytometry analysis of DAPI staining at days 3, 7, and 15, showing progressive increase in cell death after 5-FU (200μM) treatment. (B) Longitudinal brightfield and mCherry fluorescence images of the same individual organoids tracked from day 30 to day 36 post-drug withdrawal. Scale bar, 200μm. (C) Kaplan-Meier analysis of relapse-free survival (RFS) of CRC patients stratified by high or low expression of the Pdk1^high^ gene signature. (D) Kaplan-Meier analysis of post-progression survival (PPS) in the same patient cohort, stratified by high or low expression of the Pdk1^high^ gene signature. Numbers at risk are shown below the graph.

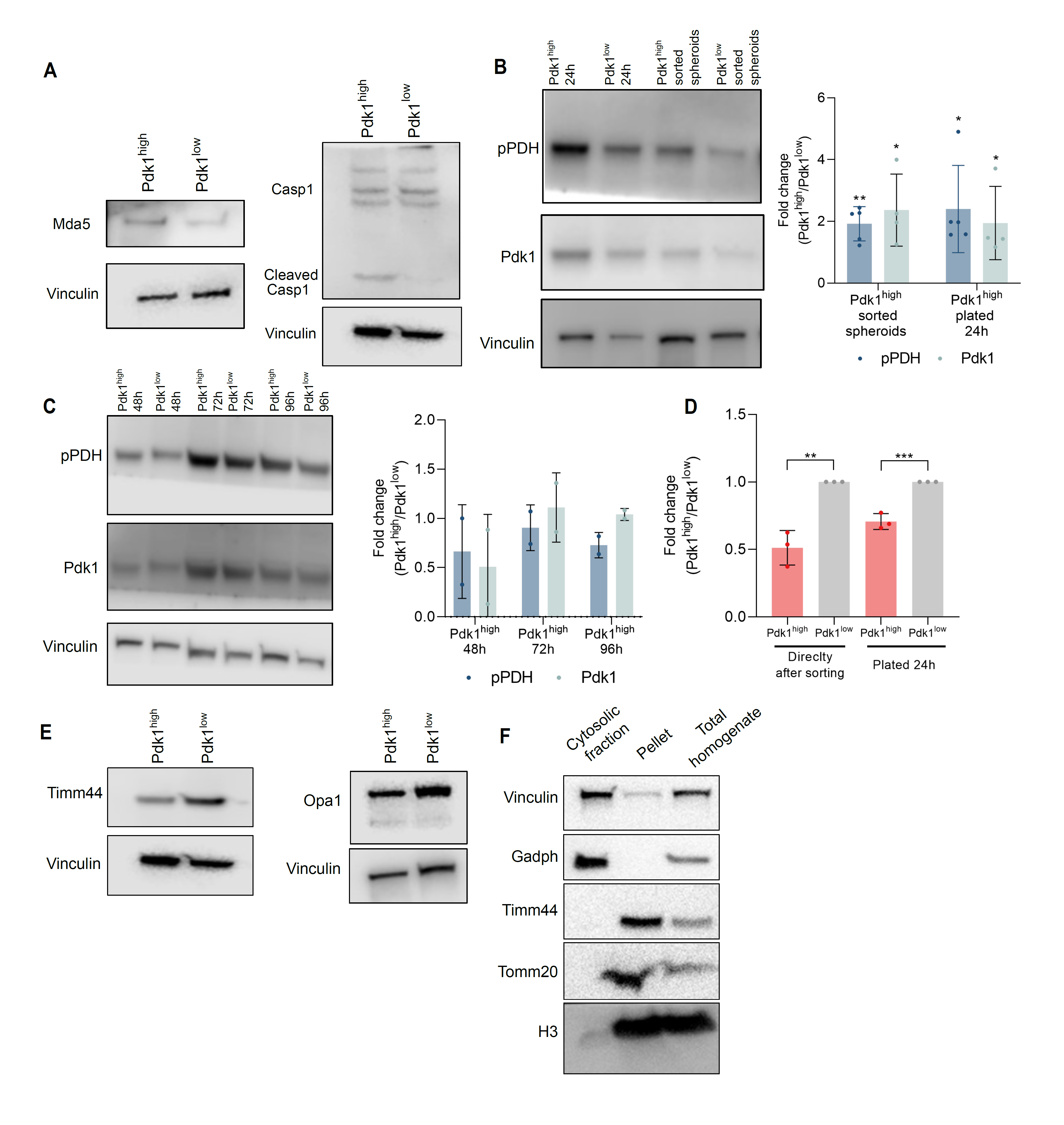

**Fig S6. Analysis of IFN- and inflammasome pathway-related protein expression in Pdk1^high^ cells.**

(A) Western blot analysis of Mda5, caspase-1, and cleaved caspase-1 protein levels in Pdk1^high^ and Pdk1^low^ cells FACS-sorted from MC38-Pdk1-mCherry spheroids. (B) Representative western blot image (left panel) and quantification (right panel) of phospho-PDH (pPDH) and PDK1 in Pdk1^high^ and Pdk1^low^ cells FACS-sorted from MC38-Pdk1-mCherry spheroids and analyzed either immediately after sorting or following 24 hours of 2D culture (n = 4–5). Data are expressed as log₂ fold change (Pdk1^high^/Pdk1^low^). Data are expressed as mean± SEM. Statistical significance was assessed using a one-sample t-test, *p < 0.05, **p < 0.01. (C) Representative western blot image (left panel) and quantification (right panel) of pPDH and PDK1 in Pdk1^high^ and Pdk1^low^ cells sorted from MC38-Pdk1-mCherry spheroids and analyzed at 48-, 72-, and 96-hours post-plating in 2D culture (n = 2). (D) Quantification of PDH activity expressed as fold change (Pdk1^high^/Pdk1^low^) in Pdk1^high^ and Pdk1^low^ cells FACS-sorted from MC38-Pdk1-mCherry spheroids, measured immediately after sorting and 24 hours after 2D replating (n = 3). Data are expressed as mean± SEM. Statistical significance was assessed using unpaired t-test, **p < 0.01, ***p < 0.001. (E) Western blot analysis of TIMM44 and OPA1 protein levels in Pdk1^high^ and Pdk1^low^ cells FACS-sorted from MC38-Pdk1-mCherry spheroids. (F) Western blot analysis of cytosolic fractionation quality in MC38 cells. Histone H3 (nuclear marker), Vinculin and GAPDH (cytosolic markers), and TIMM44 and TOM20 (mitochondrial markers) were assessed in the total homogenate, cytosolic, and pellet fractions to confirm the specificity and purity of the fractionation.

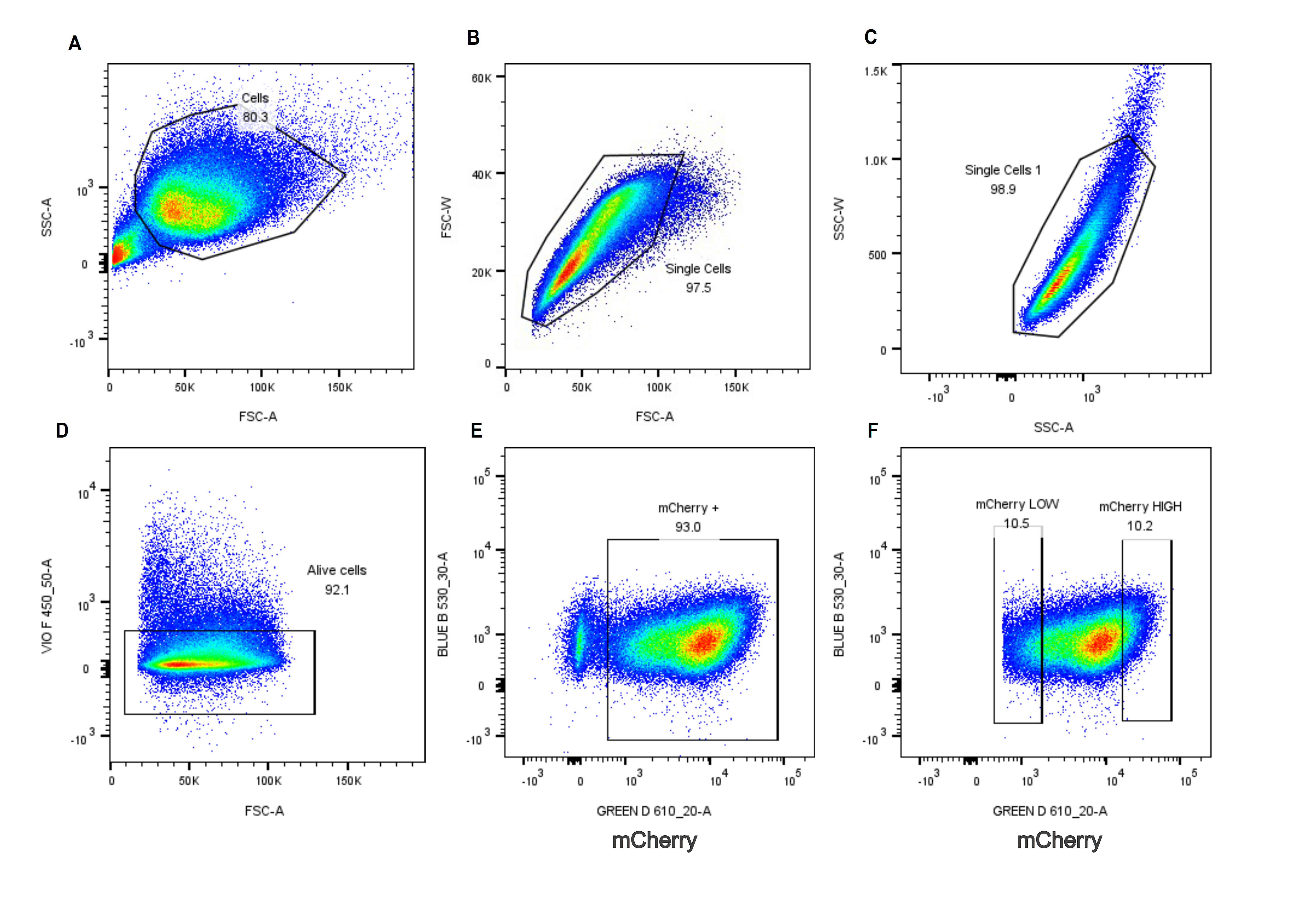

Fig S7. Gating strategy for the identification of mCherry^high^ and mCherry^low^ cells.

(A) Initial scatter gate (FSC-A vs SSC-A) used to identify the main cell population and exclude debris. (B) First singlet discrimination gate (FSC-W vs FSC-A) to exclude cell doublets and aggregates. (C) Second singlet discrimination gate (SSC-W vs SSC-A) for further exclusion of doublets. (D) Viability gate based on VIO F 450 (DAPI) vs FSC-A, used to exclude dead cells and retain viable single cells for downstream analysis. (E) Gate applied to identify mCherry-expressing cells (mCherry+), based on mCherry fluorescence intensity (GREEN D 610_20-A) vs BLUE B 530_30-A. (F) Within the mCherry+ gate, cells were further stratified into Pdk1-mCherry^low^ (bottom 10%) and Pdk1-mCherry^high^ (top 10%) subpopulations based on mCherry fluorescence intensity.
